# A chemical counterpunch: *Chromobacterium violaceum* ATCC31532 produces violacein in response to translation-inhibiting antibiotics

**DOI:** 10.1101/589192

**Authors:** Gabriel L. Lozano, Changhui Guan, Yanzhuan Cao, Bradley R. Borlee, Nichole A. Broderick, Eric V. Stabb, Jo Handelsman

## Abstract

Bacterially produced antibiotics play important roles in microbial interactions and competition. Antibiosis can induce resistance mechanisms in target organisms and may induce other countermeasures as well. Here, we show that hygromycin A from *Streptomyces* sp. 2AW induces *Chromobacterium violaceum* ATCC31532 to produce the purple antibiotic violacein. Sublethal doses of other antibiotics that similarly target the polypeptide elongation step of translation likewise induced violacein production, unlike antibiotics with different targets. *C. violaceum* biofilm formation and virulence against *Drosophila melanogaster* were also induced by translation-inhibiting antibiotics, and we identified an <u>a</u>ntibiotic-<u>i</u>nduced <u>r</u>esponse (*air*) two-component regulatory system that is required for these responses. Genetic analyses indicated a connection between the Air system, quorum-dependent signaling, and the negative regulator VioS, leading us to propose a model for induction of violacein production. This work suggests a novel mechanism of interspecies interaction in which a bacterium produces an antibiotic in response to inhibition by another bacterium.

## INTRODUCTION

In many microbial communities, diverse species contribute to complex functions that they cannot perform in isolation (Newman & Banfield, 2002). Community members also compete with each other (Ghoul & Mitri, 2016), in part through production of antibiotics—secondary metabolites that inhibit other community members. The ability for microorganisms to detect and respond to antibiotics is likely to be important for survival and competitiveness in complex communities.

Actinobacteria are prolific producers of secondary metabolites that affect development and secondary metabolism in target bacteria (Abrudan et al., 2015; Traxler & Kolter, 2015). Different classes of secreted secondary metabolites, such as siderophores, biosurfactants, and antibiotics modulate bacterial interactions. In several pathogenic bacteria, sublethal concentrations of antibiotics induce a global transcriptional response, which might be a stress response but might also indicate that antibiotics act as signal molecules (Fajardo & Martínez, 2008; Goh et al., 2002). A current challenge is to understand how bacteria transduce antibiotic exposure to a targeted transcriptional response. Evidence suggests that antibiotics typically elicit physiological responses through their inhibitory activity rather than by other means such as structural recognition (Boehm et al., 2009; Dörr et al., 2016). Cellular damage generated by bactericidal antibiotics can induce transcription of stress-response genes, but it is less clear how bacteriostatic antibiotics elicit transcriptional changes. The concept of “competition sensing” suggests that some microbes may have evolved the ability to detect a hazard signal using established stress responses and respond by up-regulating production of toxins and antibiotics (Cornforth & Foster, 2013).

*Chromobacterium* species are Gram-negative β-proteobacteria well known for production of violacein, a purple pigment with antimicrobial and anti-parasitic activities (Durán et al., 2011; Gillis & Logan, 2005). We discovered an inter-species interaction that triggers violacein production, in which *Streptomyces* sp. 2AW (Stulberg et al., 2016) induces the production of violacein in *Chromobacterium violaceum* ATCC31532. The work presented here demonstrates that production of violacein in *C. violaceum* ATCC31532 is regulated by ribosomal perturbation generated by bacteriostatic translation-inhibiting antibiotics and modulated by a previously unknown two-component regulatory system.

## RESULTS

### Hygromycin A stimulates production of violacein

We found that *Streptomyces* sp. 2AW induces the production of violacein by *C. violaceum* ATCC31532 when the bacteria are grown in close proximity (Figure 1A). Contact is not necessary, suggesting that a diffusible molecule produced by *Streptomyces* sp. 2AW is responsible for triggering the response. Partially purified hygromycin A from *Streptomyces* sp. 2AW at sublethal levels induces violacein production, as does another hygromycin A producing bacterium, *Streptomyces hygroscopicus* NRRL 2388 (Figure 1B-C). Mutations that attenuate or block hygromycin A production (Δ*hyg17 or* Δ*hyg8*, respectively (Palaniappan et al., 2009)) eliminated violacein induction by *S. hygroscopicus* (Figure 1C). Taken together these results indicate that hygromycin A is likely responsible for the ability of *Streptomyces* sp. 2AW to induce violacein production in *C. violaceum* ATCC31532.

**Figure 1.**
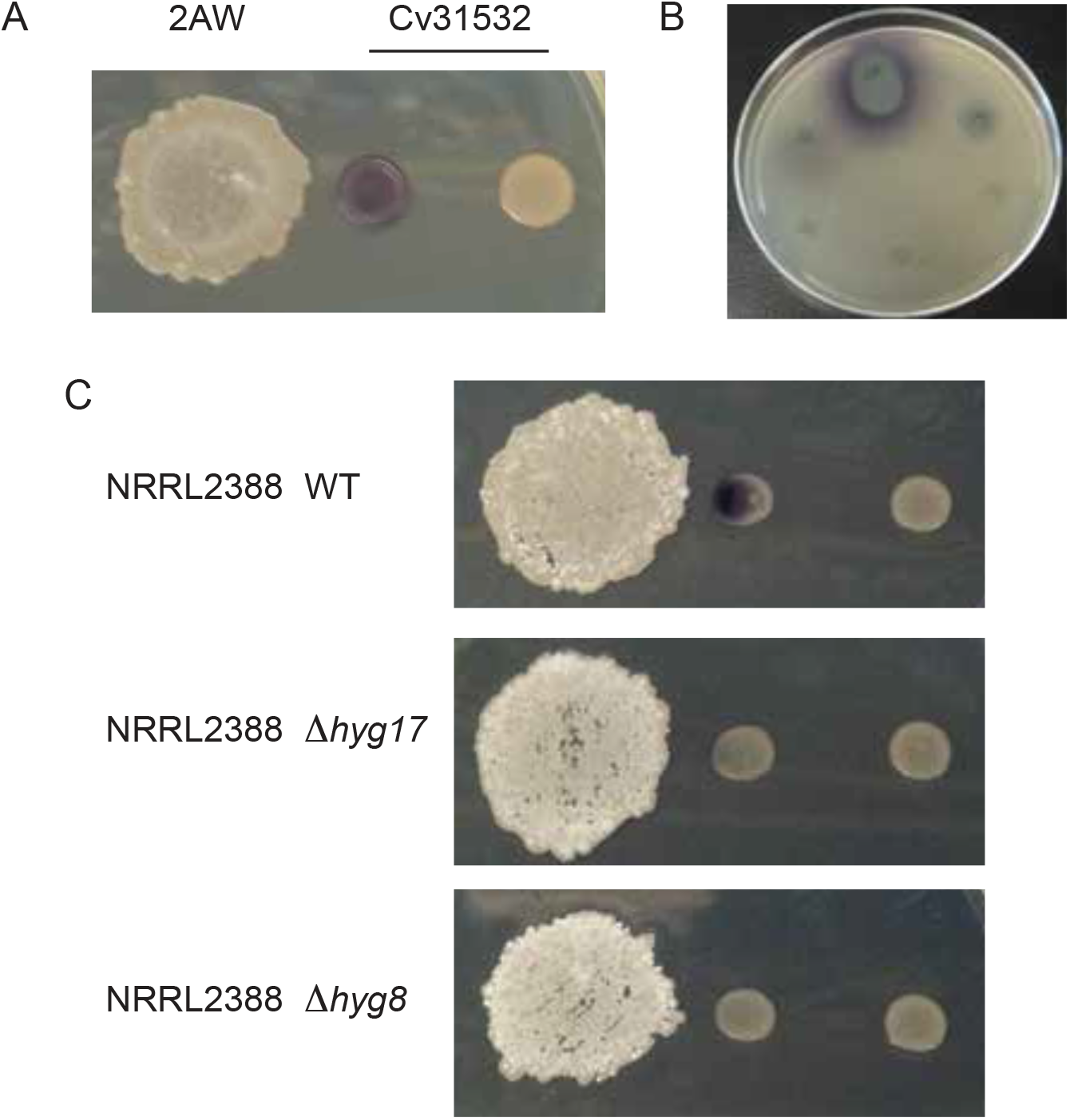
Violacein production by *C. violaceum* ATCC31532 (Cv31532) is induced by antibiotics produced by *Streptomyces* spp. (A) *C. violaceum* ATCC31532 growth with *Streptomyces* sp. 2AW (2AW). (B) HPLC fractions of methanol extract from *Streptomyces sp*. 2AW culture spotted on solid medium spread with *C. violaceum* ATCC31532. (C) *C. violaceum* ATCC31532 growth with *S. hygroscopicus* NRRL 2388 (NRRL2388) wild type (WT) and two mutants with reduced (*Δhyg17*) or abolished (*Δhyg8*) hygromycin A production.

### Violacein production is induced by inhibitors of polypeptide elongation

We considered the alternative possibilities that violacein induction could be a response to: (i) the hygromycin A molecule specifically, (ii) hygromycin A’s inhibition of translation, or (iii) sublethal antibiosis more generally. To distinguish among these three alternatives, we evaluated diverse classes of antibiotics, including those that block various steps in translation and others that have different cellular targets (Figure supplement 1). Of the twenty antibiotics tested, seven induced violacein production in *C. violaceum* ATCC31532, including blasticidin S, spectinomycin, hygromycin B, apramycin, tetracycline, erythromycin, and chloramphenicol (Figure 2A, Figure supplement 1). These antibiotics share two characteristics with hygromycin A: they inhibit growth of *C. violaceum* ATCC31532, and they block polypeptide elongation during translation, although they belong to different chemical families and inhibit translation by binding to different sites in the ribosomal region responsible for polypeptide elongation. Other antibiotics, including several that block different steps in translation (e.g., kasugamycin, puromycin, and kanamycin), did not induce violacein (Figure supplement 1).

**Figure 2.**
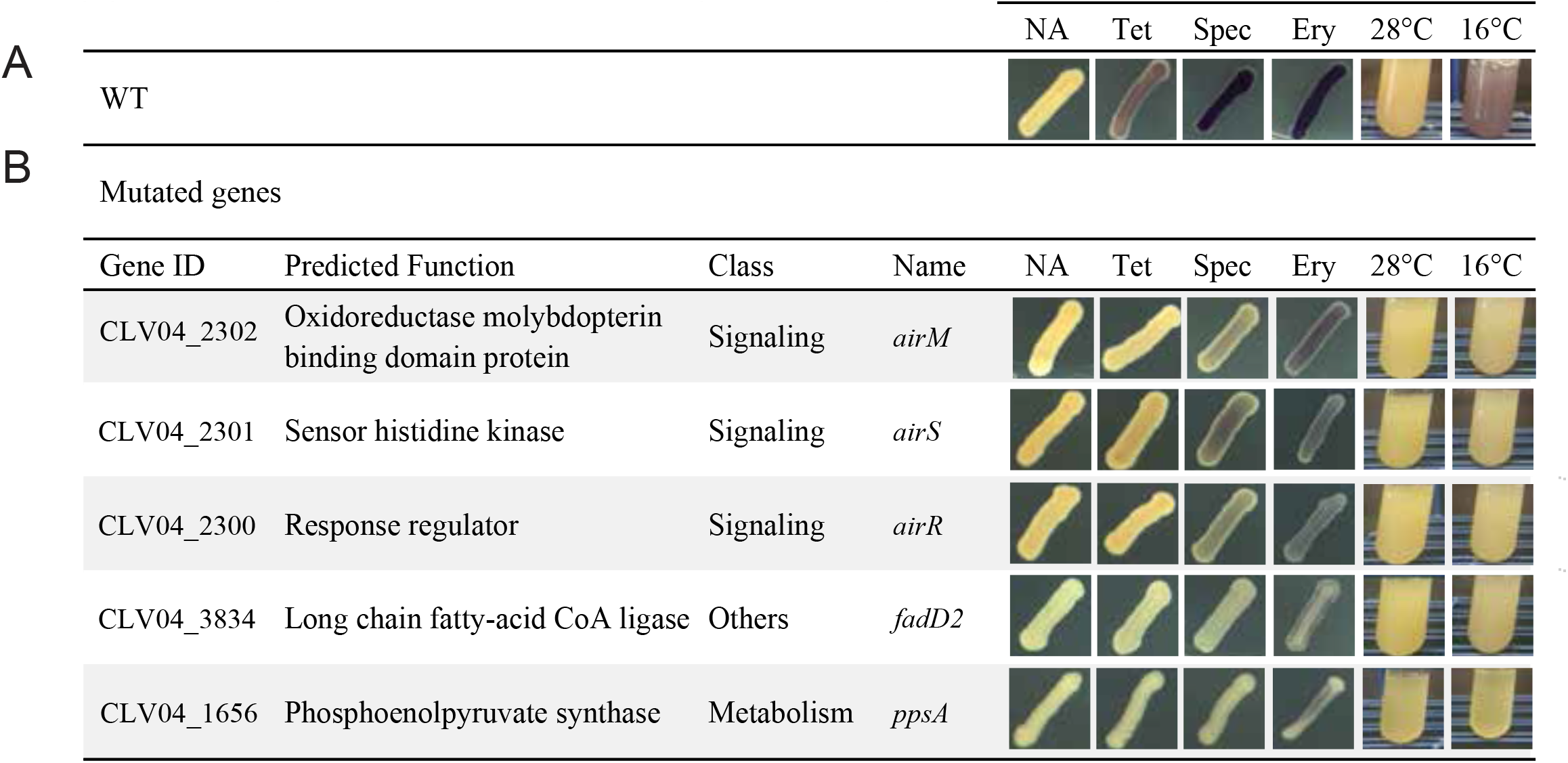
Genes involved in induction of violacein production. (A) Violacein production in response to several inducers in *C. violaceum* ATCC31532 wild type (WT). (B) *C. violaceum* ATCC31532 mutants affected in violacein production in response to all inducers tested. NA, No Antibiotic. Tet, Tetracycline. Spec, Spectinomycin. Ery, Erythromycin.

To explore whether induction of violacein production is a response to the inhibition of polypeptide elongation, we subjected *C. violaceum* ATCC31532 to cold shock. Sudden decreases in temperature can inhibit polypeptide elongation by generating secondary structures in mRNA (Horn, Hofweber, Kremer, & Kalbitzer, 2007), and previous studies indicated parallels in responses between translation-inhibiting antibiotics and cold-shock (VanBogelen & Neidhardt, 1990). We found that rapid transfer of exponential phase broth cultures from 28°C to 16°C induced violacein production in *C. violaceum* ATCC31532 (Figure 2A).

### The Air two-component regulatory system is required for the response to translation inhibition

To identify elements that participate in transducing the stimulus of translation inhibition into the response of induced violacein production, we screened random transposon mutants for loss of this ability. Because hygromycin A is not commercially available, we screened responses to sublethal concentrations of tetracycline, another strong inducer of violacein production (Figure 2A). Mutants were selected for further characterization if the screen revealed at least two independent mutants with transposon insertions in the same gene. To test the role of these genes in the regulatory response to disruption of polypeptide elongation, we further evaluated each mutant’s violacein production when treated with spectinomycin, erythromycin, or cold shock induction at 16°C. We identified five genes that, when disrupted, decrease violacein production in response to each of these stimuli (Figure 2B). We also identified mutants with altered responses to a subset of treatments (Figure supplement 2).

#### Mutants with similar responses to inhibitors of polypeptide elongation

Strains with mutations in a three-gene cluster encoding a putative two-component regulatory system do not respond to the three antibiotics tested or to cold shock (Figure 2B). We designated this cluster the <u>a</u>ntibiotic-<u>i</u>nduced <u>r</u>esponse (*air*) system, composed of genes that encode proteins predicted to serve as a sensor histidine kinase (AirS), a response regulator (AirR), and an oxidoreductase molybdopterin-binding domain (OxMoco) (IPR036374) protein (AirM). The *airS* and *airM* genes appear to be organized in an operon. In many two-component regulatory systems the sensor and response regulator genes are co-transcribed. However, in this system *airMS* and *airR* are convergently transcribed (Figure 3A-B). Notably, three other sensor histidine kinase genes in the genome are similarly arranged near genes encoding an OxMoco domain. Similar systems are also observed in other *Chromobacterium* and *Burkholderia* spp. (Figure supplement 3). To determine whether *airM* is essential for induction of violacein production, or if the phenotype of the *airM* transposon mutant reflects a polar effect on *airS*, we deleted the *airMS* operon and then complemented with *airS* or with *airMS*. Complementation with *airMS* restored the response to tetracycline whereas supplying *airS* alone did not (Figure 3C), suggesting that *airM* provides an important functional role for the two-component regulatory system.

**Figure 3.**
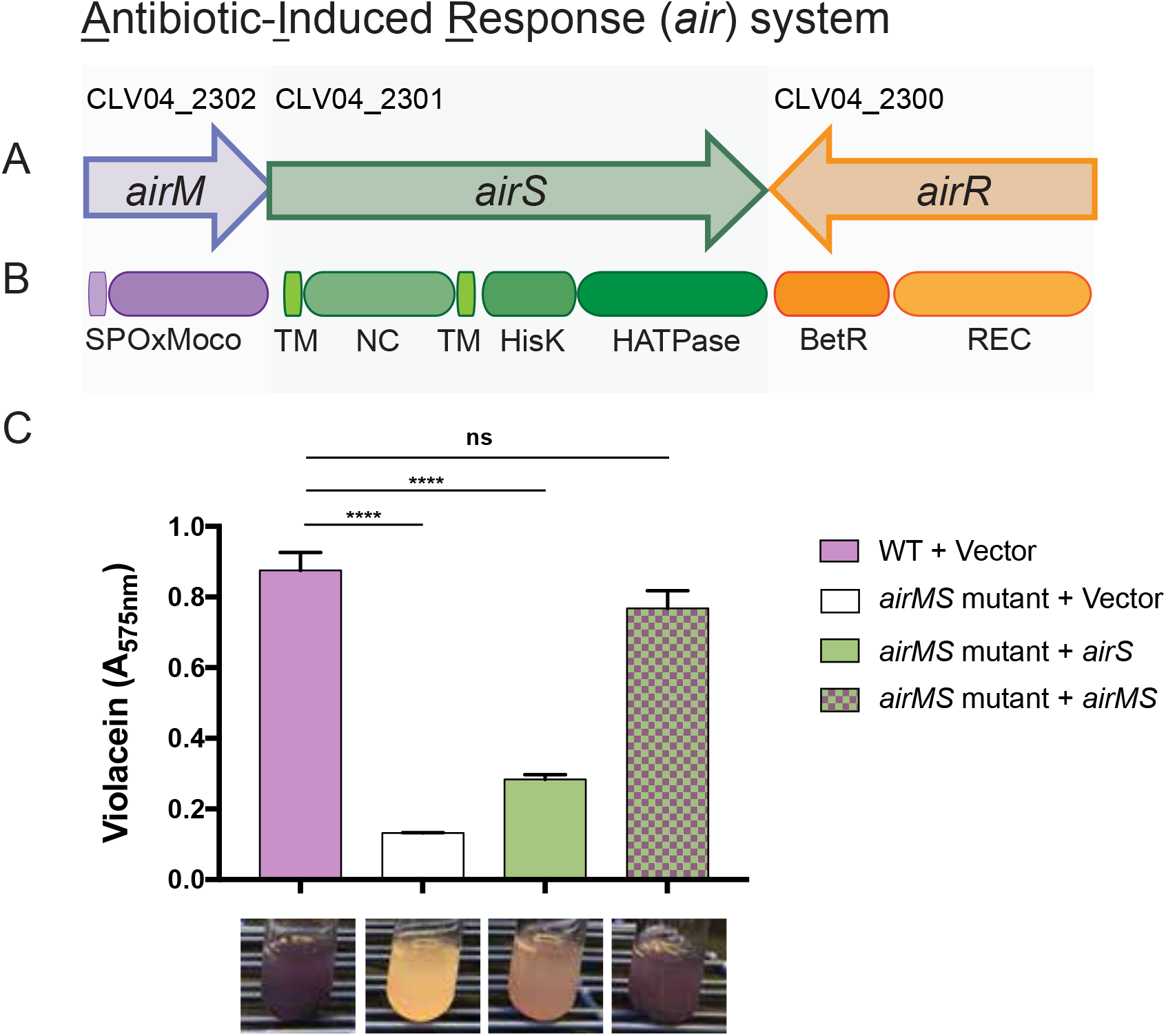
Antibiotic-induced response system. Two-component regulatory system identified by mutant analysis. (A) Gene organization; (B) Functional domains in predicted proteins. SP, signal peptide; OxMoco, oxidoreductase molybdopterin-binding domain superfamily (IPR036374); TM, transmembrane domain; NC, non-cytoplasmic domain; HisK, signal transduction histidine kinase, dimerization/phosphoacceptor domain superfamily (IPR036097); HATPase, histidine kinase/HSP90-like ATPase superfamily (IPR036890); REC, CheY-like phosphoacceptor receiver domain (IPR001789); BetR, beta-proteobacterial transcriptional regulator (IPR013975). (C) Production of violacein in wild type (WT) carrying an empty vector and in *airMS* mutant with empty vector or vector carrying *airS* or *airMS*. **** P ≤ 0.0001; ns, not significant (P > 0.05).

In addition to the mutants with insertions disrupting the *air* system, we identified strains containing transposon insertions in a phosphoenolpyruvate synthase gene (*ppsA*; CLV04_1656) and a putative long chain fatty-acid CoA ligase gene (*fadD2*; CLV04_3834) that likewise display attenuated violacein induction in response to tetracycline or other conditions that inhibit polypeptide elongation (Figure 2B).

#### Mutants with antibiotic-specific affects

We also identified several mutants that failed to induce violacein production but only for a specific subset of antibiotics (Figure supplement 2A). Upon examination, these mutants, including the recently described *cdeR* (CLV04_2412) transcriptional repressor of the *cdeAB*-*oprM* multidrug efflux pump (Evans et al., 2018), had simply become more resistant to the respective antibiotic, and at higher doses violacein induction was still evident (Figure supplement 4).

In addition, strains with mutations in a transcriptional regulator of the GntR family (CLV04_3464) and in an ABC transporter (CLV04_3178), showed a more pronounced induction of violacein in response to the presence of tetracycline, erythromycin, or spectinomycin. Strains with mutations in a putative enoyl-CoA hydratase (*fadB2*; CLV04_1011) likewise showed greater induction of violacein in the presence of the three antibiotics and to the cold shock induction at 16°C, but these mutants also had a higher basal level of violacein production even in the absence of these stimuli (Figure supplement 2B).

### Additional responses to sub-lethal concentrations of antibiotics

Sublethal concentrations of tetracycline induced other phenotypes besides violacein production in *C. violaceum* ATCC31532. For example, *C. violaceum* ATCC31532 produced biofilms on glass in response to sublethal tetracycline concentrations in an *air*-dependent manner (Figure supplement 5). We also tested *C. violaceum* ATCC31532 in an oral infection assay with *Drosophila melanogaster*, which was of interest because of the strain’s close phylogenetic association with *C. subtsugae*, an insect pathogen (Figure supplement 6). *C. violaceum* ATCC31532 killed *D. melanogaster* in the presence of tetracycline, but not in its absence, and this tetracycline-induced virulence required the *air* system (Figure 4).

**Figure 4.**
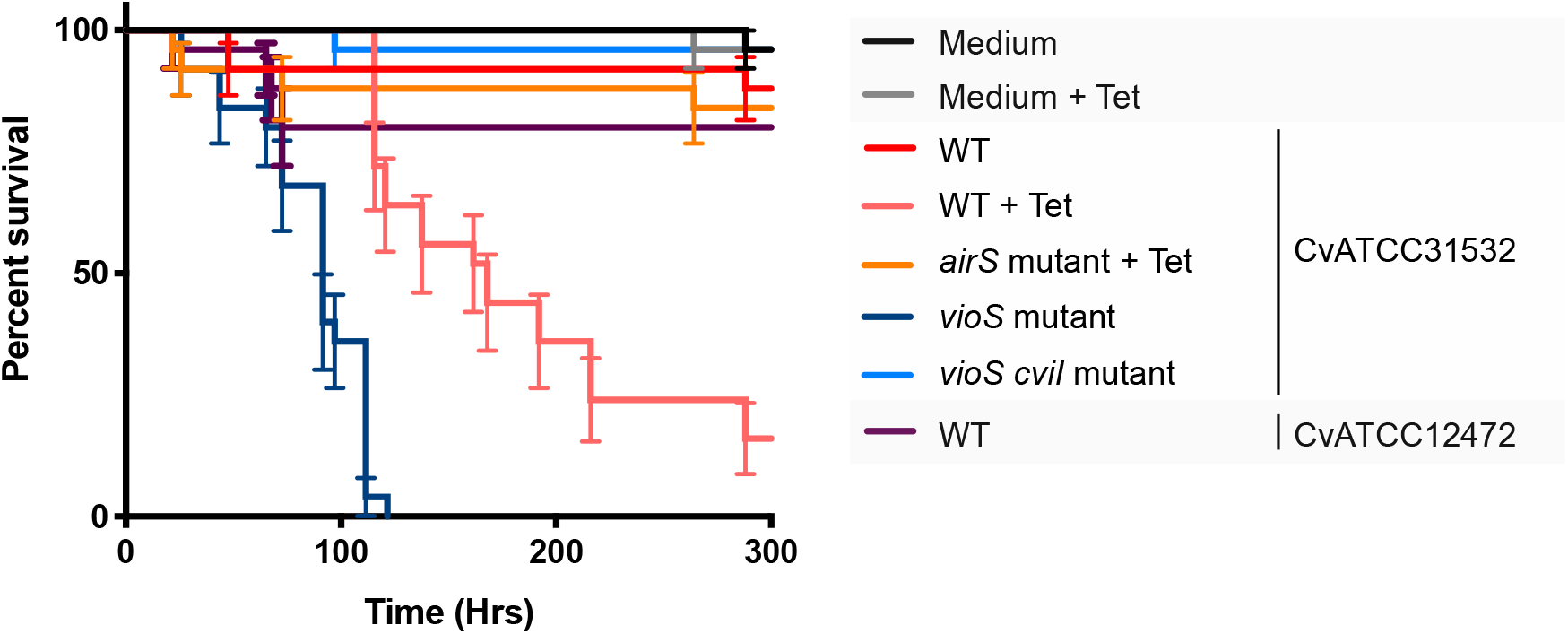
Insecticidal activity of *C. violaceum* ATCC31532 is enhanced by tetracycline. Insecticidal activity of *C. violaceum* against *Drosophila melanogaster* in response to a sublethal concentration of tetracycline. *C. violaceum* ATCC31332 (CvATCC31532) wild type (WT), *airS*, *vioS*, *vioS cviI* mutants, and *C. violaceum* ATCC12472 (CvATCC12472) wild type (WT) were evaluated. Tet, Tetracycline.

Violacein expression is under control of the CviI/CviR quorum sensing system, and it is negatively regulated by *vioS*, an otherwise uncharacterized regulator (Devescovi et al., 2017; Swem et al., 2009). We tested biofilm formation and insecticidal activity in a *vioS* mutant, which produces violacein constitutively, and in a *vioS cviI* double mutant, which does not produce violacein. Both biofilm production and insecticidal activity are expressed in the *vioS* mutant and not in the *vioS cviI* double mutant, indicating that they are regulated by the CviI/CviR quorum sensing system and repressed by VioS (Figure 4, Figure supplement 5).

### Transcriptional changes in response to sublethal concentrations of antibiotics

To understand better the physiological response to sub-lethal concentrations of antibiotics and the role of the *air* system in it, we used global RNA sequencing analysis. *C. violaceum* ATCC31532 wild type and *airR* mutant were grown both with no antibiotics and challenged separately with tetracycline and spectinomycin, and RNA pools were subjected to RNA-sequencing analysis. Each antibiotic induced a distinct but overlapping transcriptional response in the wild type (Figure supplement 7A). Using Clusters of Orthologous Groups (COG) categories, we analyzed the 640 genes that responded similarly to both antibiotics (Table supplement 1). Motility genes were enriched among genes that were down-regulated in response to tetracycline and spectinomycin (Figure supplement 7B). Genes that involved in translation, ribosomal structure and biogenesis, and secondary metabolite biosynthesis, transport, and catabolism were enriched among those up-regulated in response to both antibiotics (Figure supplement 7B).

A comparison of the WT transcriptional response with the *airR* mutant response identified 83 genes that were differentially regulated, suggesting they were directly or indirectly modulated by the *air* system. These transcripts included the violacein gene cluster and two other gene clusters encoding secondary metabolite biosynthetic pathways (Table supplement 1). Other differentially expressed genes fell in several functional categories with no distinct pattern.

Some genes that were described above as being identified in the transposon-mutant screen for altered violacein-induction responses were also found to be regulated in response to tetracycline and spectinomycin. For example, disruption of a gene that encodes a MarR family transcriptional regulator (CLV04_1869) resulted in loss of violacein induction specifically in response to tetracycline (Figure supplement 2), and this gene was upregulated in response to both spectinomycin and tetracycline (Table supplement 2). In contrast, disruption of *fadB2*, with a predicted function of an enoyl-CoA hydratase, resulted in a stronger up-regulation of violacein production but also higher background expression (Figure supplement 2), and this gene was down-regulated in response to both spectinomycin and tetracycline (Table supplement 2).

### Mechanisms of violacein induction in response to inhibitors of polypeptide elongation

Two known regulators of violacein production, *vioS* and *cviR*, were differentially expressed in the presence of either tetracycline or spectinomycin (Table supplement 1). VioS represses violacein production (Devescovi et al., 2017; McClean et al., 1997; Swem et al., 2009), and CviR is the pheromone-sensing transcriptional activator of a quorum-dependent regulatory system that activates violacein production (Stauff & Bassler, 2011; Swem et al., 2009). The RNA-seq results for these genes of interest were corroborated and expanded using targeted q-RT-PCR (Figure 5A). Transcription of *vioS* is down-regulated by sub-lethal levels of tetracycline in the wild type and in the *airR* mutant, whereas *cviR* is up-regulated in the presence of tetracycline but only in wild-type and not in the *airR* mutant (Figure 5A). The apparent requirement of *airR* for *cviR* induction was confirmed by complementing the *airR* mutant with *airR in trans* (Figure 5A). Thus, *airR* is required for the induction of *cviR* expression in response to tetracycline.

**Figure 5.**
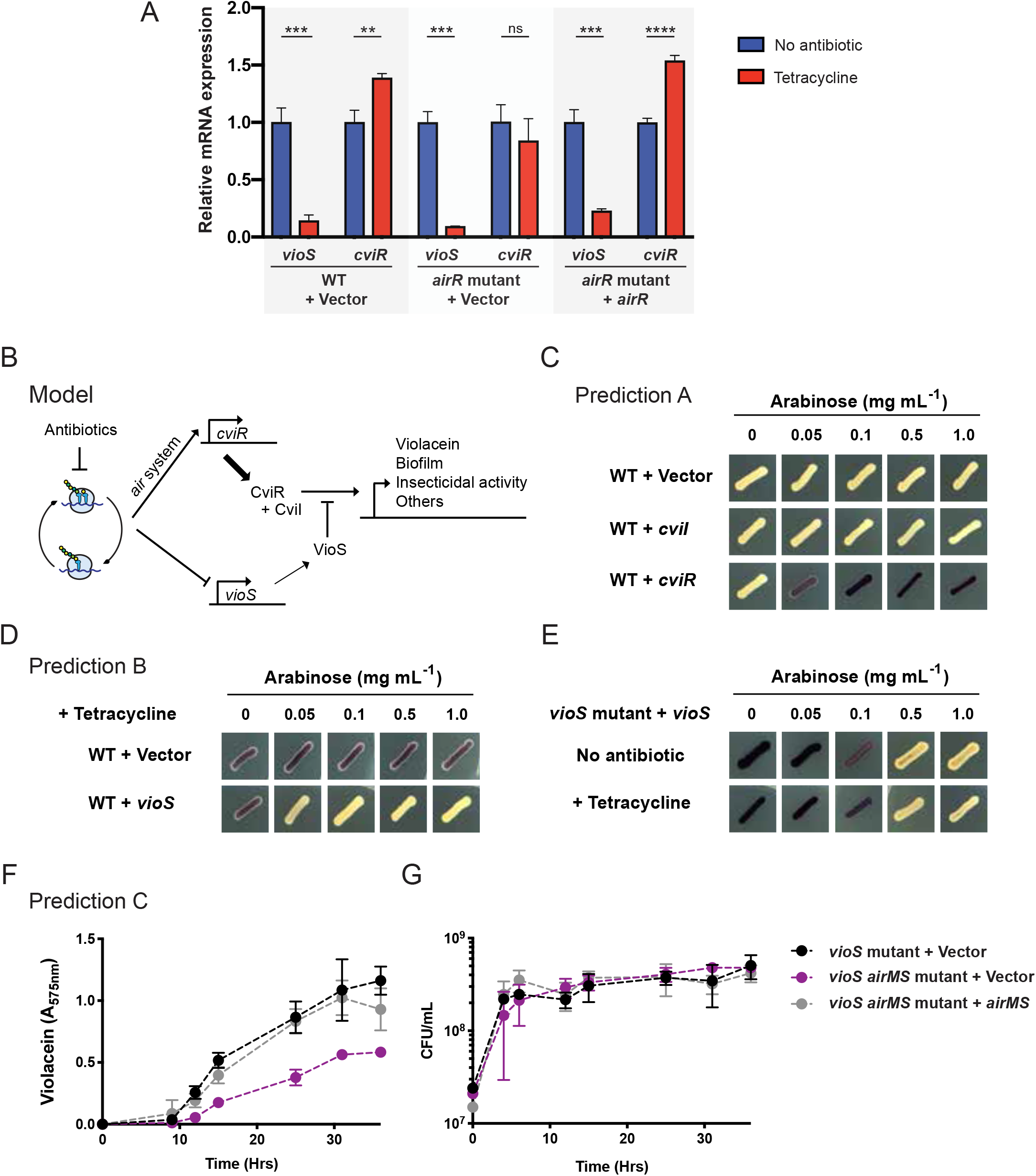
A sublethal concentration of tetracycline bypasses *vioS* repression of violacein production mediated by differential expression of *vioS* and *cviR*. (A) mRNA levels of *vioS* and *cviR* from *C. violaceum* ATCC31532 wild type (WT) carrying and empty vector, and *airR* mutant carrying an empty vector, and an *airR* mutant carrying a wild-type copy of *airR* in the presence and absence of tetracycline. ** P ≤ 0.01; **** P ≤ 0.001; **** P ≤ 0.0001; ns, no significant (P > 0.05). (B) Propose model for violacein induction by translation inhibitors. (C) Overexpression of *cviI* and *cviR* under arabinose regulation. (D) Overexpression of *vioS* under arabinose regulation in the presence of sublethal concentration of tetracycline. (E) Complementation of *vioS* mutant with *vioS* gene under regulation by arabisone, with and without tetracycline. (F) Violacein production by *vioS* and *vioS airMS* mutants. (E) Growth of *vioS* and *vioS airMS* mutants. Symbol legend applies to both (F) and (G).

Devescovi et al. recently showed that VioS is sufficient to inhibit expression of the transcriptional promoter upstream of the violacein biosynthetic gene cluster, and counteracts activation of this promoter by CviR/N-hexanoyl-L-homoserine lactone (C6-HSL) in *E. coli* (Devescovi et al., 2017). Although the mechanism of *vioS-*mediated inhibition of the *vioA* promoter is not known, the data suggest that VioS and CviR-AHL compete for the *vioA* promoter, or that VioS binds the CviR-AHL complex, thereby blocking its activity. These observations suggest that conditions favoring CviR levels over VioS levels would favor violacein production. We therefore hypothesized that the induction of violacein by sublethal concentration of antibiotics resulted from two independent mechanisms, decreased *vioS* expression and increased *cviR* expression mediated by the *air* system (Figure 5B).

We drew three predictions from this model (Figure 5B): (A) increasing the levels of the violacein activator CviR would bypass repression by VioS; (B) constitutive overexpression of *vioS* would block violacein induction by translation-inhibiting antibiotics; and (C) the *air* system would mediate the violacein-induction response to translation-inhibiting antibiotics even in the absence of *vioS*. We validated predictions A and B, as follows. Overexpression of *cviR* induces violacein with no antibiotics added, while overexpression of *cviI*, the quorum sensing autoinducer synthase, does not induce violacein production (Figure 5C). This observation suggests that under these conditions, the quorum-dependent response is limited by CviR levels but not by the autoinducer levels. Also, constitutive overexpression of *vioS* blocks violacein induction in the presence of translation-inhibiting antibiotics (Figure 5D-E).

To test prediction C, we first generated the *vioS airMS* double mutant. Unexpectedly, this double mutant produced less violacein than the *vioS* single mutant without tetracycline (Figure 5F), and without impacting growth fitness (Figure 5G). The results suggested some activity of the Air system even in the absence of tetracycline. Consistent with this possibility, comparison between the wild type and *airR* mutant RNA-seq data in the absence of antibiotics showed that the *air* system affected regulation of at least 15 genes (Table supplement 3). Thus, the *air* system appears to have some activity even without a translation-inhibition signal. Importantly, the data in Figure 5F show that the *air* system modulates violacein production independently of VioS as we predicted above.

## DISCUSSION

In this study, we examined an inter-bacterial interaction mediated by sublethal levels of antibiotics. We found that *C. violaceum* ATCC31532 produces violacein in response to sublethal levels of hygromycin A released from *Streptomyces* sp. 2AW, and in response to other structurally diverse bacteriostatic antibiotics that inhibit the elongation step of translation. Genetic analysis in *C. violaceum* ATCC31532 revealed a newly described two-component regulatory complex, the *air* system, that participates in the regulation of violacein production as well as virulence and biofilm production, all of which are regulated by the CviI/CviR quorum sensing system. Transcriptomic analysis of the wild type and *airR* mutant showed antibiotic-mediated down-regulation of *vioS* and up-regulation of *cviR*, revealing a mechanism in which VioS repression of violacein is overcome. This new inter-bacterial competition mechanism differs from the previously identified competition strategy of acyl-homoserine lactone-dependent eavesdropping (Chandler, Heilmann, Mittler, & Greenberg, 2012), and suggest that *C. violaceum* ATCC31532 can sense and respond to other members of the microbial community in part by using transcriptional regulators that detect inhibitory effects of secondary metabolites produced by their neighbors.

The idea that antibiotics serve as signals in microbial communities (Fajardo & Martínez, 2008) is supported by our findings that sublethal levels of hygromycin A produced by *Streptomyces sp.* 2AW induce violacein production by *C. violaceum* ATCC31532 when the bacteria grow in close proximity. Further experiments are required to determine whether hygromycin A plays a signaling role in natural communities. As observed in human pathogenic bacteria, sub-lethal concentrations of antibiotics in *C. violaceum* ATCC31532 also influence social behavior such as pathogenesis, biofilm formation, quorum sensing, and secondary metabolite production (Andersson & Hughes, 2014). In those systems, it appears that antibiotics function as signals through cellular damage caused by the inducing antibiotic and detected by general stress response networks (Cornforth & Foster, 2013). *C. violaceum* ATCC31532 produces violacein only in response to inhibitors of the polypeptide elongation step of translation. Recently, Liu et al. reported a similar phenomenon in which translation-inhibitors induce sliding motility in *Bacillus subtilis* (Y. Liu, Kyle, & Straight, 2018). *C. violaceum* ATCC31532 also produces violacein in response to cold shock, providing another example of the long-known parallel between responses to translation-inhibiting antibiotics and cold-shock (VanBogelen & Neidhardt, 1990). These findings suggest that the activity of the antibiotics, in this case inhibition of the polypeptide elongation step of translation, creates a cellular stress that initiates a signaling cascade.

The *air* system consists of a two-component regulatory system, a sensor (AirS) and a response regulator (AirR), and an oxidoreductase molybdopterin-binding protein (AirM), an unexpected element based on prototypical two-component regulators. In a broad database analysis, we found *airM*-like genes associated with two-component regulatory systems mainly in β-proteobacteria, but the *air* system is the first identified with an associated function. The *air* system is puzzling because a predicted membrane sensor, AirS, detects a cytoplasmic perturbation of ribosome activity. We hypothesize that the perturbation of actively translating ribosomes would create several cellular changes detected by the *air* system. We cannot infer the nature of the signal detected by AirS, since there is no annotation of known sensor domains in the AirS sequence. The predicted function of AirM, an oxidoreductase protein, suggests that the signal might involve oxidative change. This is compatible with the observed up-regulation of the NADH ubiquinone oxidoreductase complex, although expression of oxidative stress pathways did not appear to change (data not shown).

Another potential signal could be alteration in the lipid composition of the membrane. Our genetic screen identified two genes, a long chain fatty-acid CoA ligase (*fadB2*) and an enoyl-CoA hydratase (*fadD2*), that are homologs of genes that participate in fatty acid catabolism (Fujita, Matsuoka, & Hirooka, 2007). Enoyl-CoA hydratase is down-regulated by sublethal concentrations of tetracycline and spectinomycin, and the loss-of-function mutant produces violacein without exposure to antibiotics. The long chain fatty-acid CoA might scavenge phospholipids associated with the membrane, and the down-regulation of the enoyl-CoA hydratase mediated by antibiotics could change the pool of saturated and unsaturated acyls-CoA, thereby altering the composition of the new phospholipids added to the membrane.

The *air* system is needed for maximum violacein production without sublethal concentration of tetracycline in a mutant lacking the negative regulator *vioS*, indicating that the system is active without antibiotic stress and may have a housekeeping function. This is supported by the differential gene expression of several genes mediated by the *air* system without antibiotics. In addition, the presence of a third element in this two-component system may indicate that the *air* system integrates multiple signals of different cellular pathways, as has been shown in other two-component regulatory systems with auxiliary elements (Buelow & Raivio, 2010). We hypothesized that *C. violaceum* ATCC31532 co-opts a preexisting signaling network to integrate a response generated by the ribosome perturbation. This could expand the model of “competition sensing”, whereby bacteria adapt not only to general stress response networks, but also transcriptional modulators, such as two-component regulatory systems that respond to any physiological response generated by antibiotics. The ribosome is one of the most common targets for antibiotics (Wilson, 2014), and being able to sense inhibition of its function rather than detecting each antibiotic independently might enable *C. violaceum* ATCC31532 to have a single response to many competitors rather than separate responses to individual species.

Our discovery of *C. violaceum’s* antibiotic production in response to antibiosis was facilitated by the fact that this “chemical counterpunch” (i.e., violacein) is purple. With this fortuity in mind, a central question arising from the current study is whether similar phenomena are widespread but less visible. It seems likely that microbial communities possess less obvious but equally important emergent forms of chemical competition. New approaches and technologies (Chodkowski & Shade, 2017; Harn, Powers, Shank, & Jojic, 2015) have poised the field for a more comprehensive approach to discovering chemically mediated responses underpin microbial interactions.

## ACKNOWLEDGMENTS

We gratefully acknowledge Bailey Kleven for her technical assistance, and Dr. Sara Knaack for discussing transcriptomic analysis. This research was supported by the Office of the Provost at Yale University, and by funding from the Wisconsin Alumni Research Foundation through the University of Wisconsin–Madison Office of the Vice Chancellor for Research and Graduate Education. EVS was supported by grant MCB-1716232 from the National Science Foundation.

## MATERIALS AND METHODS

### Bacterial strains and culture conditions

*Streptomyces sp.* 2AW (Schloss et al., 2010), *Streptomyces hygroscopicus* NRRL2388 (Palaniappan et al., 2009), *Chromobacterium violaceum* ATCC31532 WT, *C. violaceum* ATCC31532 *vioS* (Cv017)(McClean et al., 1997)*, C. violaceum* ATCC31532 *vioS cviI* (Cv026)(McClean et al., 1997) and *C. violaceum* ATCC12472 were cultured in LB (10 g L^−1^ tryptone; 5 g L^−1^ yeast extract; 10 g L^−1^ NaCl). *S. hygroscopicus* NRRL 2388 and the corresponding mutants were a gift from Kevin Reynolds at Portland State University. Antibiotics were obtained from Sigma (St. Louis, MO, USA) (ceftazidime, chloramphenicol, erythromycin, fusidic acid, hygromycin B, nalidixic acid, paromomycin, piperacillin, polymyxin B, puromycin, tetracycline, trimethoprim, vancomycin); from RPI (Mt Prospect IL, USA) (apramycin, blasticidin S, rifampicin, spectinomycin); from American Bio (Natick, MA, USA) (kanamycin); from MP Biomedicals (Santa Ana, CA, USA) (streptomycin); and from Enzo Life Sciences (Farmingdale, NY, USA) (kasugamycin).

### Inter-species interaction assay

*Streptomyces sp.* 2AW and *S. hygroscopicus* NRRL2388 were spotted on LB plates and incubated for 3-5 d when 5-10 μL of *C. violaceum* ATCC31532 liquid culture grown for 16 h at 28°C were spotted on two different positions on the plates. Plates were incubated at 28°C until violacein production in *C. violaceum* ATCC31232 was observed.

### Violacein induction assay

Fractions of a methanol extract from *Streptomyces sp*. 2AW grown on solid media (Stulberg et al., 2016) were directly tested against *C. violaceum* ATCC31532. Each fraction was spotted on LB plates, and then 100 μL of *C. violaceum* ATCC31532 liquid cultures grown to an OD600 ∼ 4.0 at 28°C were spread over the plates. Plates were incubated for two days at 28°C. The following antibiotics were evaluated by directly spotting 10 μL of stock solution on LB plates: apramycin (100 μg mL^−1^), blasticidin S (25 μg mL^−1^), ceftazidime (20 μg mL^−1^), chloramphenicol (34 μg mL^−1^), erythromycin (50 μg mL^−1^), fusidic acid (10 μg mL^−1^), hygromycin B (50 μg mL^−1^), kanamycin (50 μg mL^−1^), kasugamycin (10 μg mL^−1^), nalidixic acid (10 μg mL^−1^), paromomycin (10 μg mL^−1^), piperacillin (50 μg mL^−1^), polymyxin B (50 μg mL^−1^), puromycin (25 μg mL^−1^), rifampicin (20 μg mL^−1^), spectinomycin (50 μg mL^−1^), streptomycin (100 μg mL^−1^), tetracycline (10 μg mL^−1^), trimethoprim (5 μg mL^−1^), and vancomycin (10 μg mL^−1^).

### *C. violaceum* ATCC31532 transposon mutagenesis and genetic screen for mutants defective in violacein production

pSAM_BT21 was generated from pSAM_BT20 (Sivakumar et al., 2019) by interchanging the ampicillin-resistance gene with a kanamycin resistance cassette amplified from pENTR/D-TOPO using primers KanTopo_MluIFor/KanTopo_MluIRev (Table supplement 4), and inserted in the MluI site. *C. violaceum* ATCC31532 and *E. coli* S17-1λpir with pSAM_BT21 with kanamycin (50 μg mL^−1^) were first grown individually for 16 h at 28°C and 37°C, respectively, with agitation. Cells were washed and resuspended in fresh medium to an OD600 of 2.0. One volume of *E. coli* S17-1λpir with pSAM_BT21 was mixed with two volumes of *C. violaceum* ATCC31532. Cells were harvested (6,000 × *g*, 6 min), resuspended in 100 µL of fresh medium, and spotted on LB plates. The conjugation mixture was incubated at 28°C for 6 h, then scraped and resuspended in 2.5 mL of LB. 100 µL aliquots were plated on LB containing gentamicin (50 μg mL^−1^), ampicillin (200 μg mL^−1^) and tetracycline (0.125 μg mL^−1^) for selection of *C. violaceum* ATCC31532 transconjugants defective in violacein production. Plates were incubated for two days at 28°C.

For each mutant, 1 mL of liquid culture grown for 16 h was harvested (6,000 × *g*, 6 min), and cells were resuspended in 400 μL of TE (10 μM Tris HCl pH 7.4; 1 µM EDTA pH 8.0). Samples were boiled for 6 min, centrifuged (6,000 × *g*, 6 min), and 2 μL of supernatant was used as a template for DNA amplification. Transposon locations were determined by arbitrarily primed PCR (Goodman et al., 2009), which consisted of a nested PCR using first-round primer GenPATseq1 and either AR1A or AR1B and second round primer GenPATseq2 and AR2 (Table supplement 4). PCR products of the second round were purified by gel extraction (QIAquick Gel Extraction Kit; QIAGEN) and then sequenced using primer GenPATseq2. PCR sequencing was performed by the DNA Analysis Facility on Science Hill at Yale University.

### Assay for violacein production in response to antibiotics and cold shock

Violacein production by mutants identified as defective for violacein induction in response to tetracycline (0.125 μg mL^−1^) was evaluated in the presence of spectinomycin (2 μg mL^−1^), erythromycin (2 μg mL^−1^), and in liquid cultures at 16°C and 28°C. Dose-dependent response of violacein production to tetracycline (0.25 – 4 μg mL^−1^), spectinomycin (4 – 64 μg mL^−1^), and erythromycin (2 – 16 μg mL^−1^) was evaluated in several mutants on LB plates. The response to these antibiotics and cold shock was evaluated visually based on the purple color of violacein.

### Chromosomal deletion of the *airMS* operon

The *airMS* operon was deleted by allelic exchange and replaced with a chloramphenicol-resistance cassette. The *airMS* deletion cassette was constructed by a modified version of overlap extension (OE) PCR (Ho, Hunt, Horton, Pullen, & Pease, 1989). Fragments 1 kb upstream and 1 kb downstream of the *airMS* operon were amplified using primers MuCv0535/6_Afor and MuCv0535/6_Arev, or MuCv0535/6_Bfor and MuCv0535/6_Brev, respectively (Table supplement 4). The resulting products had overlapping homology and further amplification with primers MuCv0535/6_Afor and MuCv0535/6_Brev resulted in a single combined product of approximately 2 kb representing a fusion of the upstream and downstream sequences. This PCR product was cloned into pENTR/D-TOPO, generating pairMS_ENTR. Primers MuCv0535/6_Arev and MuCv0535/6_Bfor were designed to introduce a SphI site in the overlapping region to allow introduction of a selectable resistance gene. A chloramphenicol-resistance cassette was amplified from pACYC184 using primers pACYC184Cm_For/pACYC184Cm_Rev, which contain SphI sites in the 5’ region, and cloned into pENTR/D-TOPO, generating pCm_ENTR. The chloramphenicol cassette was recovered from pCm_ENTR using SphI, and cloned between the upstream and downstream region of the *airMS* operon. A *mob* element was recovered from pmob_ENTR (Lozano et al., 2019) using AscI, and cloned into an AscI site in the pENTR backbone, generating pairMS_Cm_mob_ENTR. Conjugation mixtures of *C. violaceum* ATCC31532 and *E. coli* S17-1λpir carrying the pairMS_Cm_mob_ENTR vector were prepared following the procedure for generating transposon mutants. Double recombinant *C. violaceum* ATCC31532 transconjugants were selected based on their ability to grow on chloramphenicol (34 µg mL^−1^) and screened for the inability to grow on kanamycin (50 µg mL^−1^). The *airMS* deletion mutant was confirmed by PCR using MuCv0535/6_Afor and MuCv0535/6_Brev, and by evaluating violacein production in the presence of tetracycline. The same methodology was used to delete *airMS* in the *C. violaceum* ATCC31532 *vioS* mutant.

### Complementation and overexpression assays

The broad-host-range expression and arabinose-inducible vector pJN105 was modified by introduction of the chloramphenicol-resistance cassette recovered from pCm_ENTR into the SphI site, generating pJN105Cm. *airS* was amplified using primers CV0536_For and CV0536_Rev. *airMS* was amplified using primers CV0535/6_For and CV0536_Rev. *airR* was amplified using primers CviR_For and CviR_Rev. *vioS* was amplified using primers CV1055_For and CV1055_Rev. *cviI* was amplified using primers CviI_For and CviI_Rev, and *cviR* was amplified using primers CviR_For and CviR_Rev. For all these genes, a XbaI site in the 5’ region was added to the forward primer, and a SacI site in the 5’ region was added to the reverse primer, for a directional integration of each gene in front of the *araBAD* promoter in pJN105Cm. Plasmids were transferred to the corresponding host using the same conjugation protocol used for generating the transposon mutants. Genes under control of the *araBAD* promoter were induced with arabinose (0.05 – 1 mg mL^−1^).

### Drosophila melanogaster oral infection assay

*Canton-S* (*Cs*) flies (wolbachia free) were used as standard wild-type lines. *Drosophila* stocks were maintained at 25°C on cornmeal medium (8 g L^−1^ agar; 80 g L^−1^ polenta; 40 g L^−1^ yeast; 40 g L^−1^ sucrose; 53.6 ml L^−1^ moldex). *C. violaceum* strains were streaked from frozen glycerol stocks onto LB plates and incubated at 28°C overnight. Isolated colonies were then inoculated into LB medium and cultured at 28°C for 20 h. Cultures were centrifuged (4,000 *rpm*, 20 min, 4°C). The supernatant was decanted, the pellets were resuspended in the remaining liquid, and the concentration of the cultures were adjusted to OD_600_ = 200 (approximately 100X concentration of the original overnight culture). For antibiotic treatment, tetracycline (2.5 µg mL^−1^) was added to concentrated cultures immediately before feeding to flies.

Adult female flies were starved in empty vials for 2 h at 29°C. Paper filters were placed on top of food medium and 150 μL of a 1:1 mixture of the concentrated pellet (OD600 = 200) and 2.5% sucrose was added; for the sucrose-negative control, LB was substituted for the bacterial pellet. The starved flies were transferred into the infection vials and kept at 29°C. Survival was assessed at 2 h post-infection to account for any infection-independent mortality. The number of dead flies per vial was recorded twice per day for approximately five days after infection. Potential differences in survival between treatments were analyzed for significance with Kaplan Meier Survival Analysis using Graphpad Prism software.

### Phylogenetic analysis

*Chromobacterium* spp. genomes were recovered from the NCBI database, June 2017 (Table supplement 5). Phylogenomic reconstruction was accomplished using the phylogenetic and molecular evolutionary (PhaME) analysis software (Ahmed, Lo, Li, Davenport, & Chain, 2015). PhaME identified SNPs from the core genome alignments, and the phylogenetic relationships were inferred by maximum likelihood using FastTree.

### RNAseq analysis

*C. violaceum* ATCC31532 WT and *airR* mutant were grown without antibiotics, with tetracycline (0.125 µg mL^−1^) or with spectinomycin (2 µg mL^−1^) in 5 mL of LB at 28°C with agitation in duplicate. RNA samples were prepared from 250 µL of cells grown to an OD600 ∼ 3.2. Cultures were mixed with 750 µL of TRIzol and incubated at 65°C for 10 min. Samples were frozen at −80°C for 10 min and thawed at room temperature for 15 min. Chloroform (200 µL) was added, the samples were shaken and incubated at room temperature for 3 min and centrifuged (12,000 × *g*, 15 min, 4°C). The aqueous phase was recovered, mixed with 500 µL of isopropanol, incubated at room temperature for 10 min, and centrifuged (12,000 × *g*, 10 min, 4°C). The pellet was washed with 1 mL of 75% ethanol, and air-dried for 10 minutes, resuspended with 50 µL of RNase-free water, and finally incubated at 60°C for 15 min. DNA was removed from 5 µg of total RNA using the TURBO DNase kit (Invitrogen, Carlsbad, CA, USA).

RNA samples were treated with Ribo-Zero rRNA removal kit (Illumina, San Diego, CA, USA), cDNA libraries were constructed with an average size between 150 bp and 200 bp, and they were sequenced in Illumina HiSeq2500 paired-end 2×75 platform. Library preparation and sequencing were performed by the Yale Center for Genome Analysis. Low quality sequences were trimmed using Trimmomatic (Bolger, Lohse, & Usadel, 2014). Mapped reads and estimated gene expression levels were calculated using RSEM with a transcript list of ORFs recovered from *C. violaceum* ATCC31532 genome (GenBank assembly accession GCA_002865685.1) (B. Li & Dewey, 2011). Differential expression was assessed using limma (Ritchie et al., 2015) using “voom” function to model the mean-variance relationship from read counts converted to log2-counts-per-million (logCPM). Genes that were not appreciably expressed (transcripts per million (TPM) < 5) were discarded, as recommended (Ritchie et al., 2015). Genes with an adjusted P value < 0.01 were identified as being differentially expressed.

Genes were identified as having a generalized response to inhibition of translational elongation if they were differentially expressed in WT in the presence of tetracycline and in the presence of spectinomycin, relative to no antibiotic (Table supplement 1). Genes for which the response to translational inhibition is mediated, directly or indirectly, by the *air* system were identified if they were not differentially expressed in response to tetracycline or streptomycin in the *airR* mutant background (but were in WT), or if these genes are differentially expressed when compare WT against *airR* in the presence of both antibiotics (Table supplement 6).

### Quantitative RT-PCR

Quantitative reverse-transcriptase PCR was used to validate the differential gene expression detected for *vioS* and *cviR* in the RNA-Seq analysis. Primers used are listed in Table supplement 4. *C. violaceum* ATCC31532 WT pJN105Cm, *airR* pJN105Cm, and *airR* pJN105Cm_airR were grown with chloramphenicol (34 µg mL^−1^), and with or without tetracycline (0.125 µg mL^−1^) in triplicate as reported above. Total RNA was recovered and DNAase treated in the same manner as the RNA recovered for RNA-seq analysis. Two hundred nanograms of DNase-treated RNA was reverse transcribed into cDNA using SuperScript™ III First-Strand Synthesis System (Invitrogen). Quantitative PCR was carried out in a 10-μL volume using PowerUp SYBR Green Master Mix (Applied biosystems) with 1 μL of cDNA and 200 nM PCR primers. These reactions were performed using the CFX96 Real-Time System (Bio-Rad) using the following cycling parameters: 50°C for 10 min, 95°C for 5 min, followed by 40 cycles of 95°C for 10 sec and 60°C for 30 sec. Reverse transcriptase-minus (RT-minus) template PCRs were included as negative controls to confirm the absence of genomic DNA contamination. Tenfold serially diluted DNA standard curves were included on every plate. Melting curve analysis were done to verify the specificity of the PCR products. Expression levels under each condition were normalized to the *dnaG* housekeeping gene, and the Pfaffl method was used to calculate fold change in gene expression (Pfaffl, 2001). Differences between groups were tested for statistical significance (Student’s t-test) using GraphPadPrism 7 software.

### Characterization of C. violaceum ATCC31532 vioS airR

*C. violaceum* ATCC31532 *vioS* pJN105Cm, *vioS airMS* pJN105Cm, and *vioS airMS* pJN105Cm_airMS were grown with chloramphenicol (34 µg mL^−1^) and arabinose (0.2 mg mL^−1^), with agitation for two days at 28°C. Samples were withdrawn periodically to evaluate bacterial growth by serial dilution and plating in LB, and to quantify violacein production. Violacein was quantified by a crude violacein extraction (Z. Liu et al., 2013). One-mL aliquots of cultures of each strain were centrifuged (14,000 × *g*, 20 min), and cells were resuspended in 1 mL of ethanol to dissolve violacein. Supernatants were recovered after centrifugation (12,000 × *g*, 10 min), and transferred to 96-well plates. Violacein concentration was determined spectrophotometrically at 575 nm in a Synergy HT plate reader (BioTek).

---

**Figure supplement 1.**
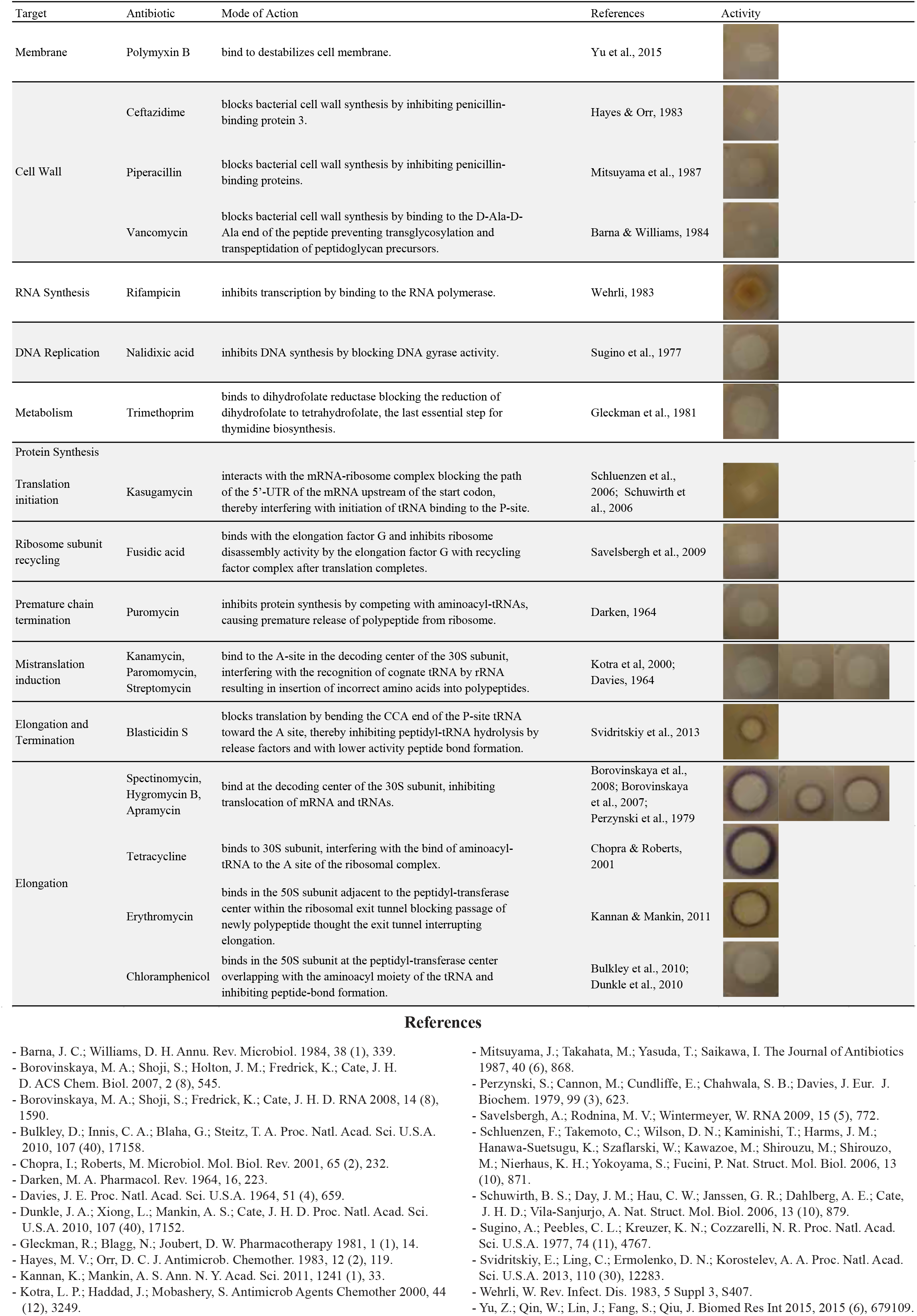
Profile of *C. violaceum* ATCC31532 violacein production in response to structurally diverse antibiotics. Antibiotics are classified by cellular target.

**Figure supplement 2.**
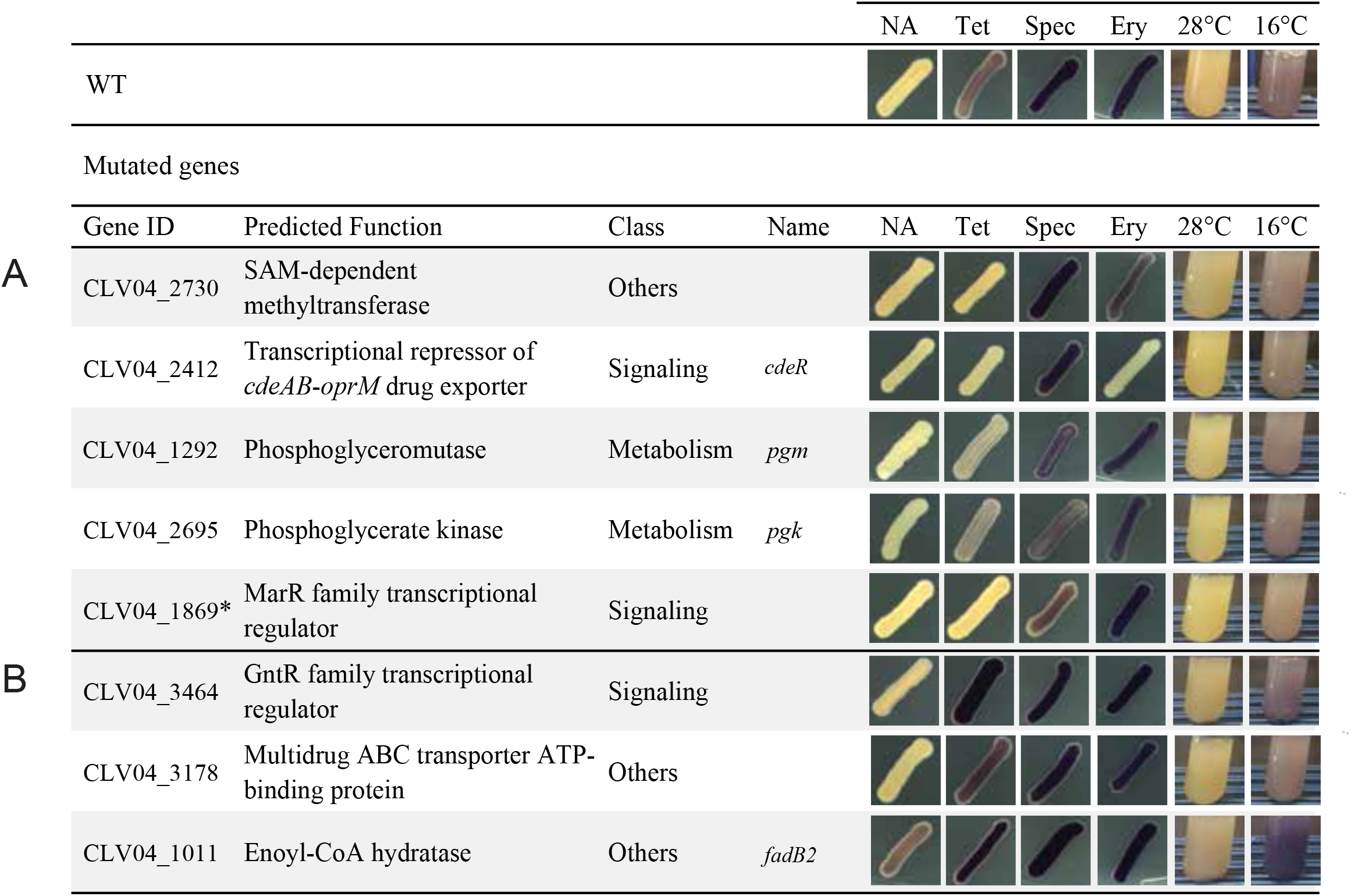
Additional genes involved in induction of violacein production by several inducers. (A) *C. violaceum* ATCC31532 mutants with a disrupted violacein production in response to some of the inducers tested. (B) *C. violaceum* ATCC31532 mutants with increased violacein production in response to some of the inducers tested. NA, No Antibiotic. Tet, Tetracycline. Spec, Spectinomycin. Ery, Erythromycin. * Transposons in promotor.

**Figure supplement 3.**
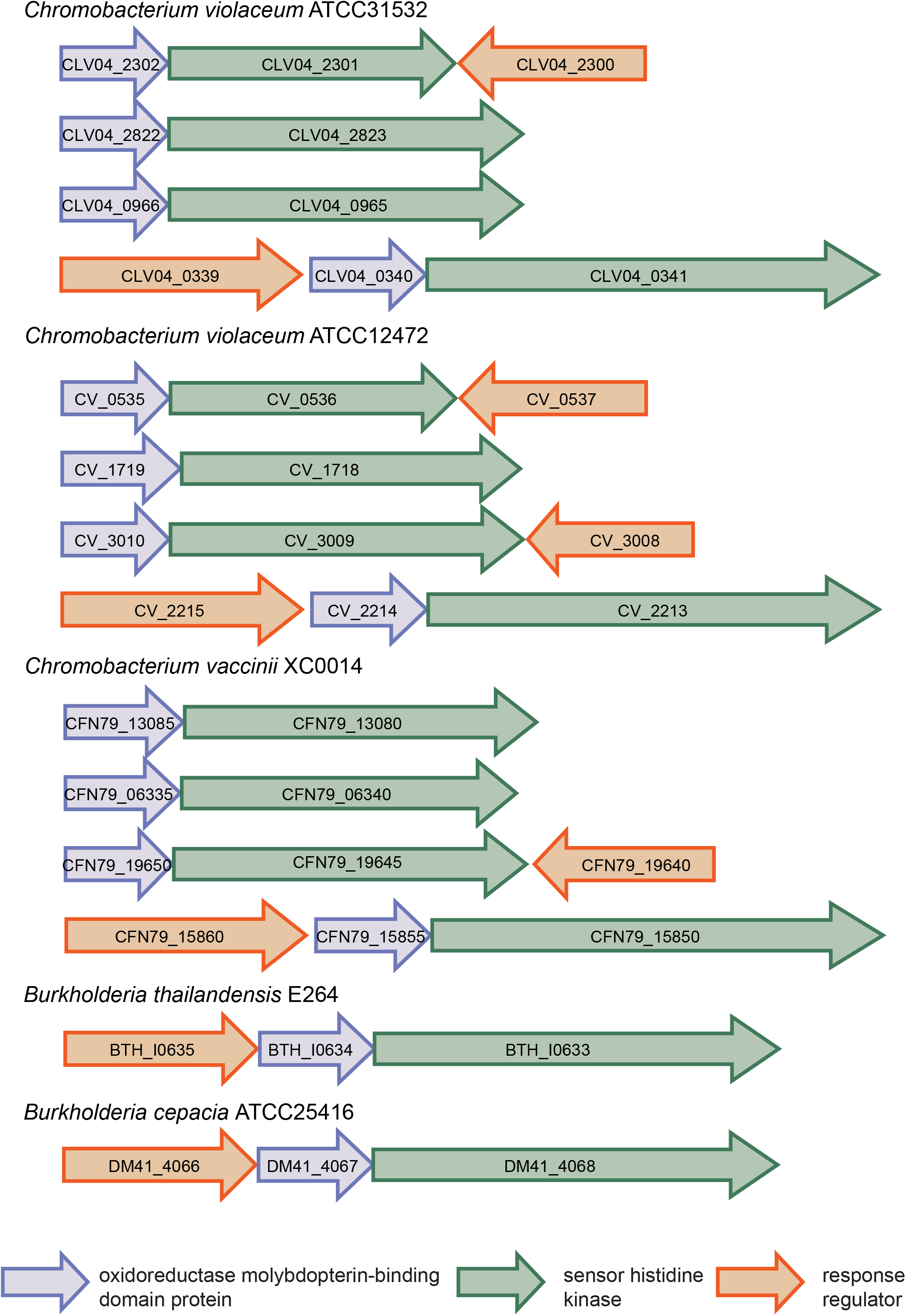
Other pair genes that encode oxidoreductase molybdopterin-binding domain (OxMoco)(IPR036374) proteins next to a sensor histidine kinases in *Chromobacterium* and *Burkholderia* spp genomes.

**Figure supplement 4.**
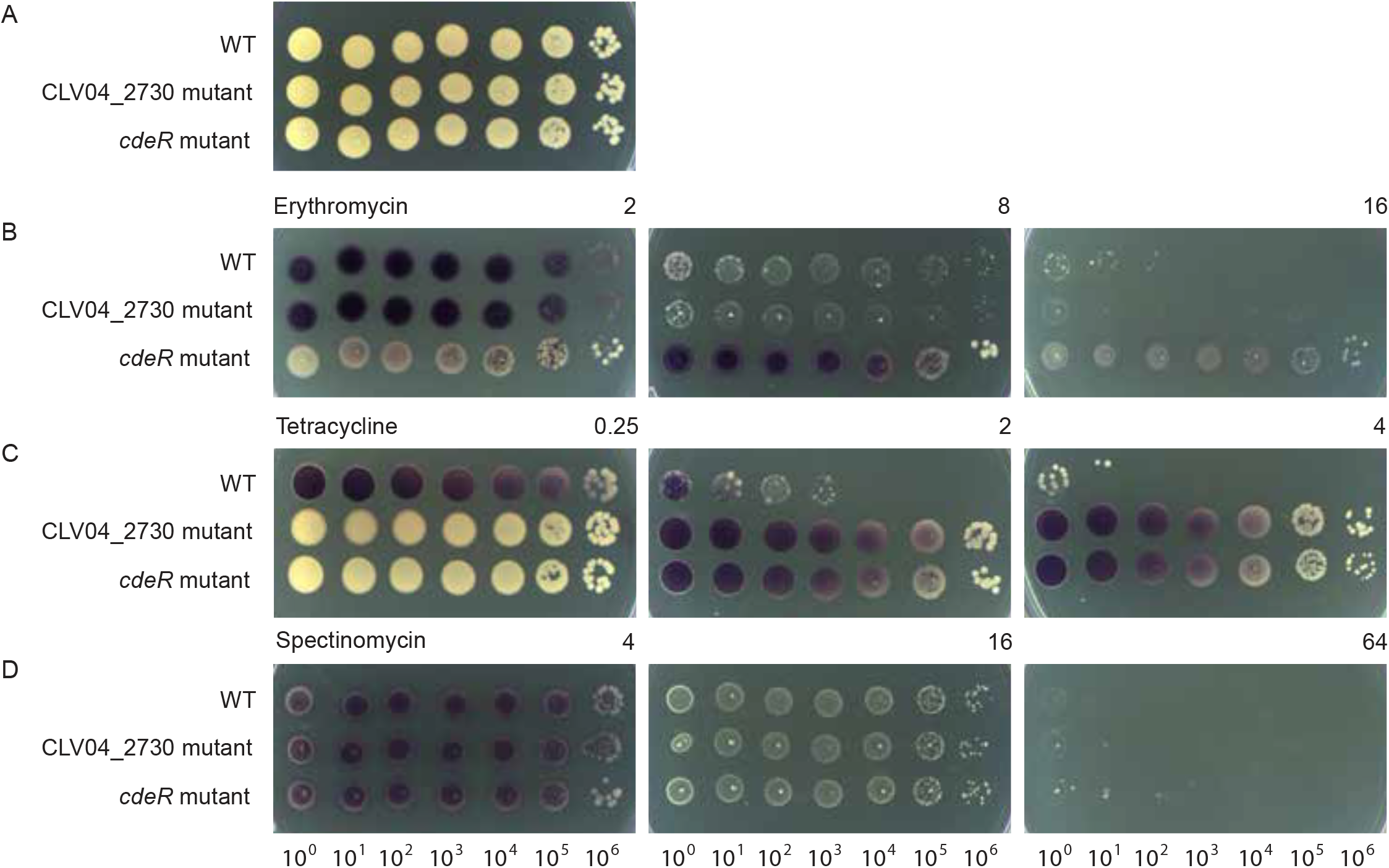
Dose-dependent violacein production in two antibiotic-resistant mutants. Survival and violacein production of *C. violaceum* ATCC31532 wild type (WT), mutant in a SAM-dependent methyltransferase (CLV04_2730), and mutant in *cdeR*, a transcriptional repressor of the *cdeAB*-*oprM* multidrug efflux pump system (CLV04_2412) with (A) no antibiotic; (B) Erythromycin (µg mL−1); (C) Tetracycline (µg mL−1); (D) Spectinomycin (µg.mL−1).

**Figure supplement 5.**
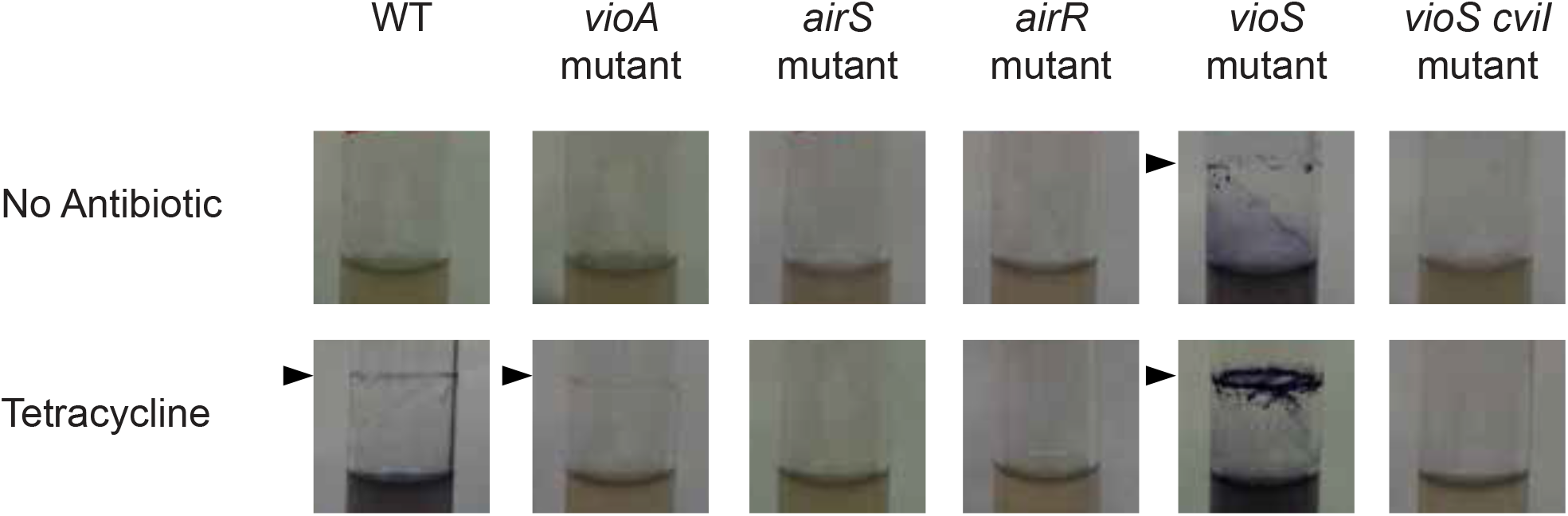
*C. violaceum* ATCC31532 biofilm formation on glass in response to a sublethal concentration of tetracycline. Biofilm formation by *C. violaceum* ATCC31532 wild type (WT), *vioA*, *airS*, *airR*, *vioS*, and *vioS cviI* mutants without antibiotics, or in the presence of tetracycline (0.25 µg.mL−1). Violacein mutant, *vioA*, was used as a control for a biofilm visualization without violacein pigmentation. Arrow indicates the location of the biofilm.

**Figure supplement 6.**
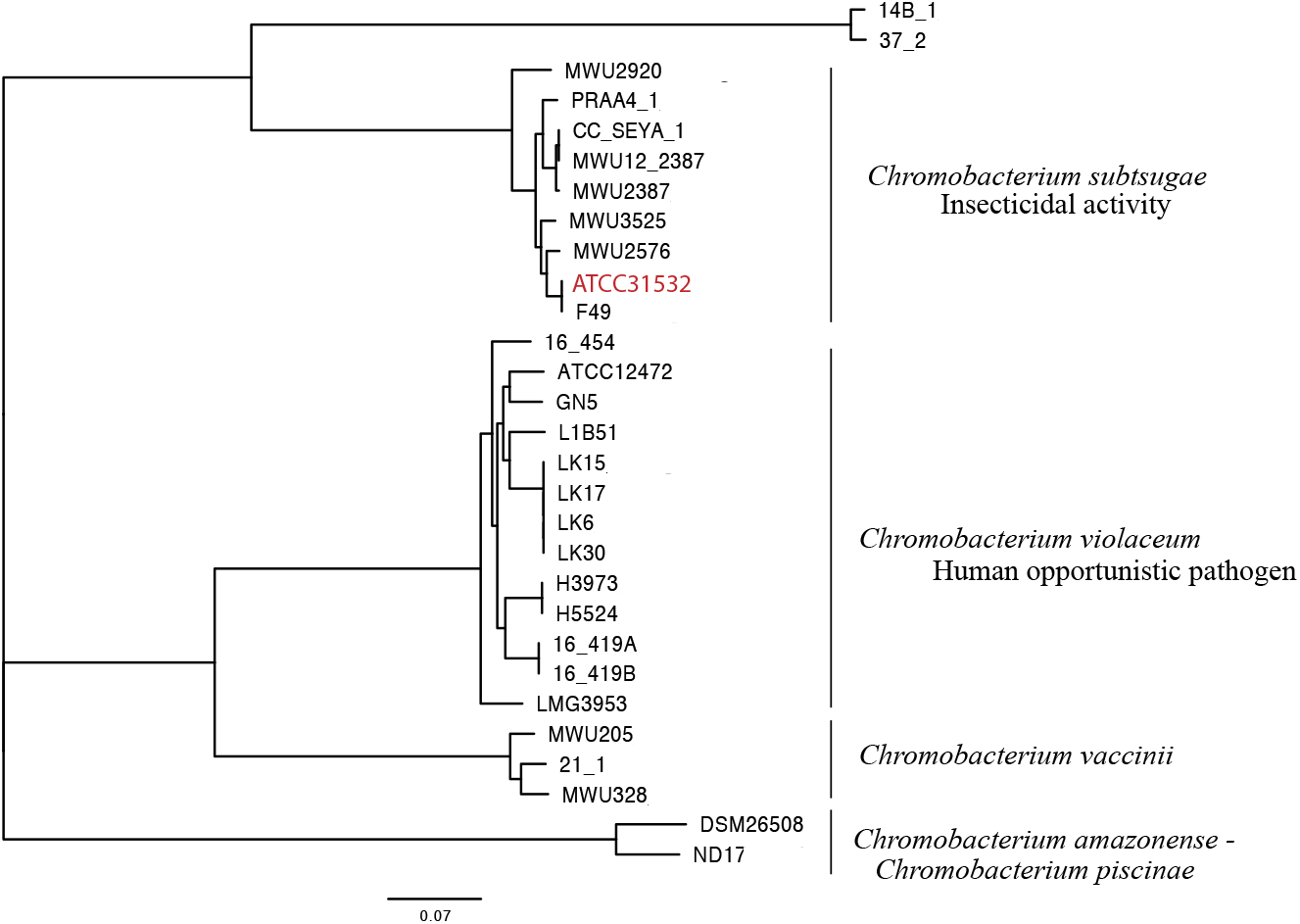
Phylogenetic relatedness of several *Chromobacterium* spp. Highlight in red *C. violaceum* ATCC31532.

**Figure supplement 7.**
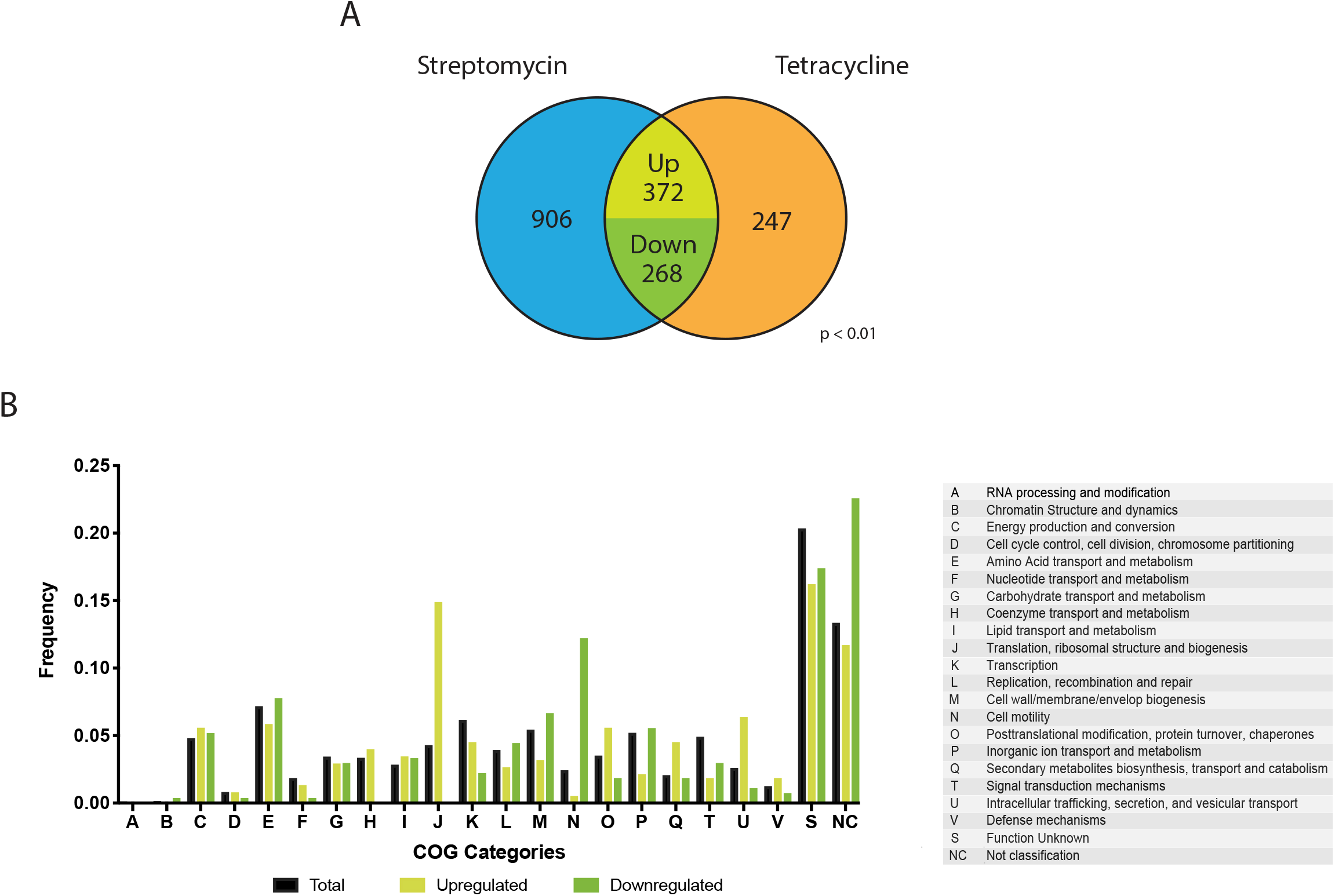
Transcriptome analysis of *C. violaceum* ATCC31532 in the presence and absence of a sublethal concentration of translation inhibitors. The analysis indicates change in expression from the baseline condition with no antibiotics. (A) Venn diagram of gene differentially expressed by *C. violaceum* ATCC31532 wild type (WT) in streptomycin or tetracycline. (B) COG classification of the differentially genes expressed in streptomycin and tetracycline.

**Table supplement 1.**
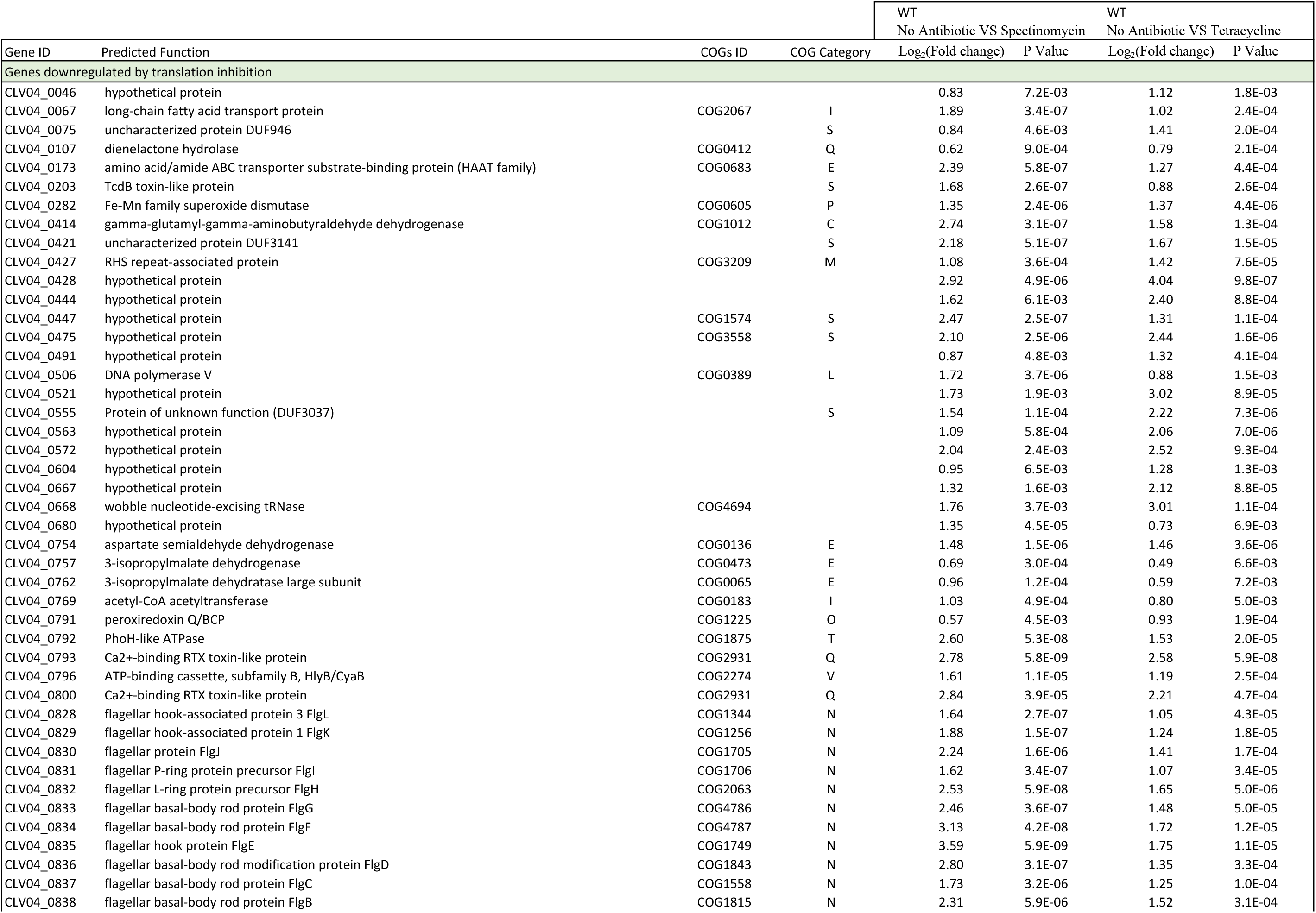
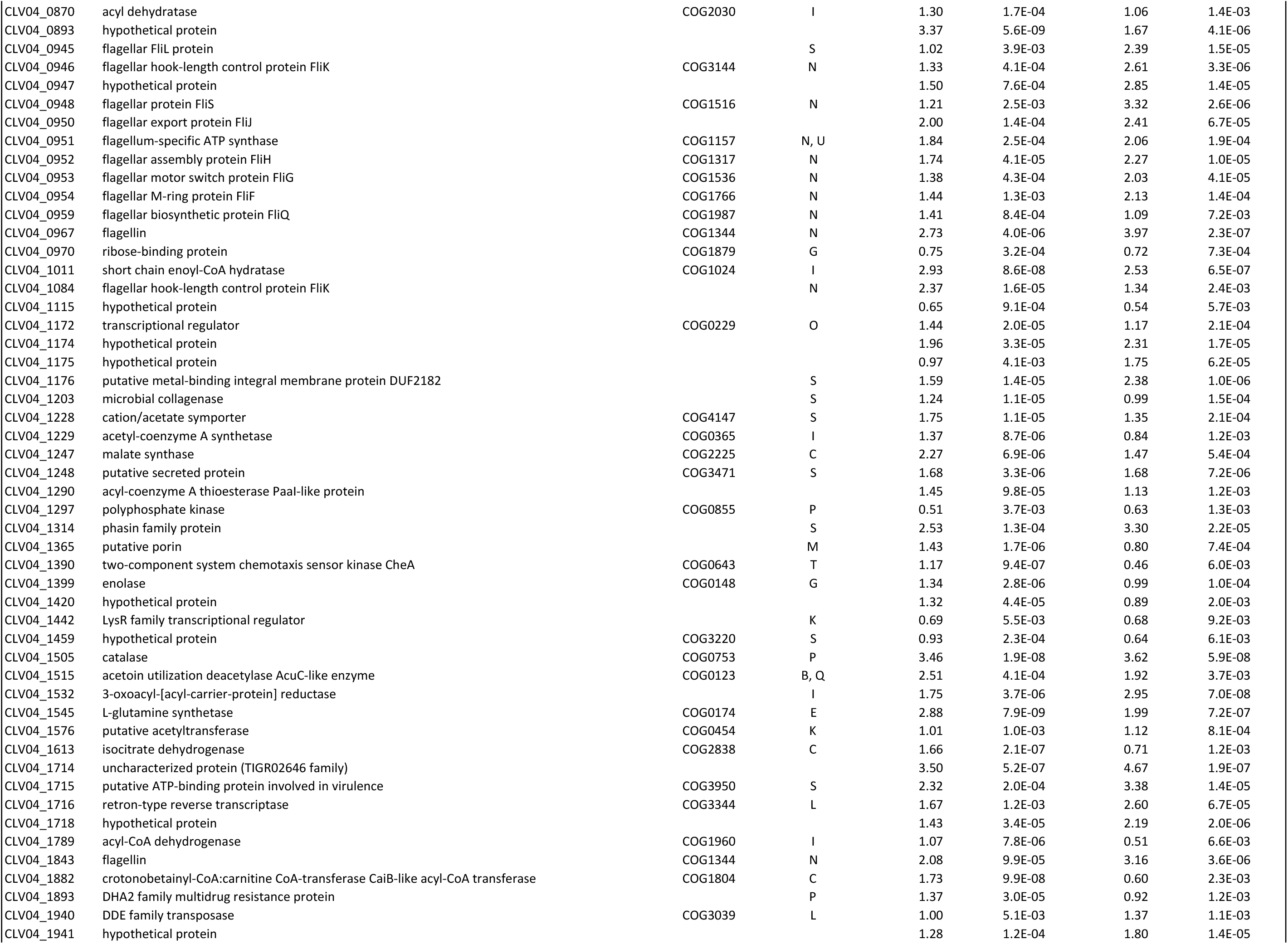
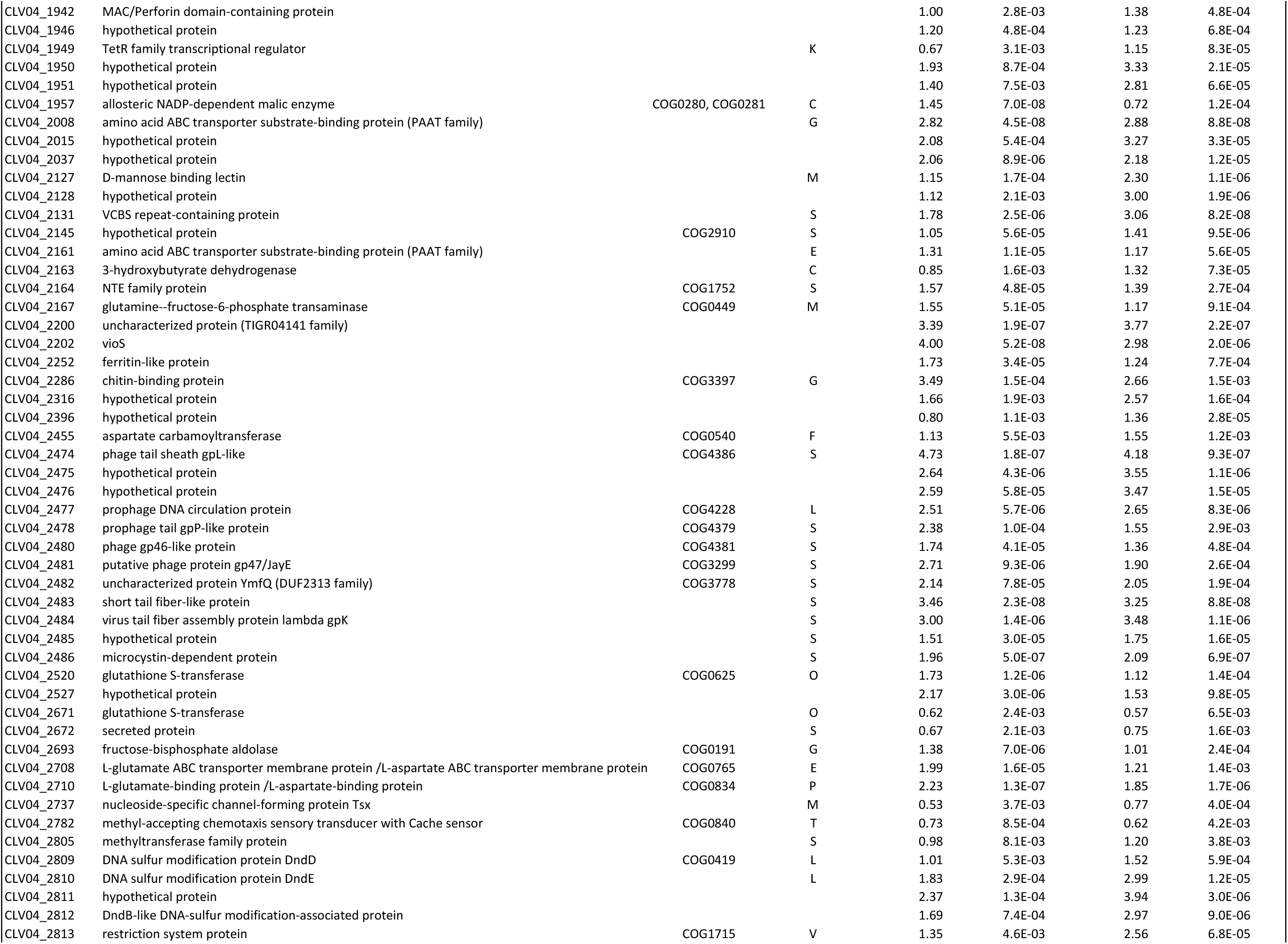
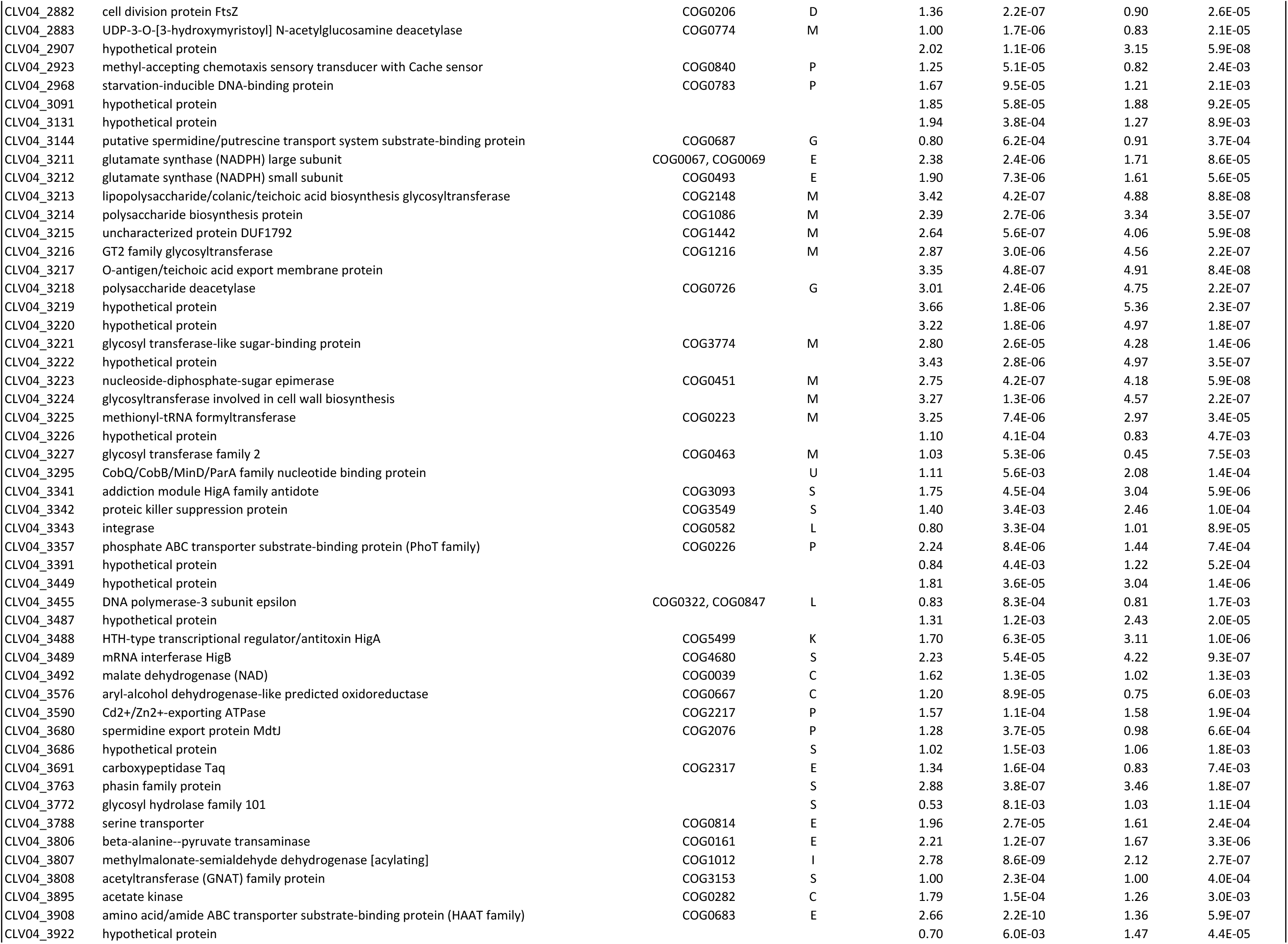
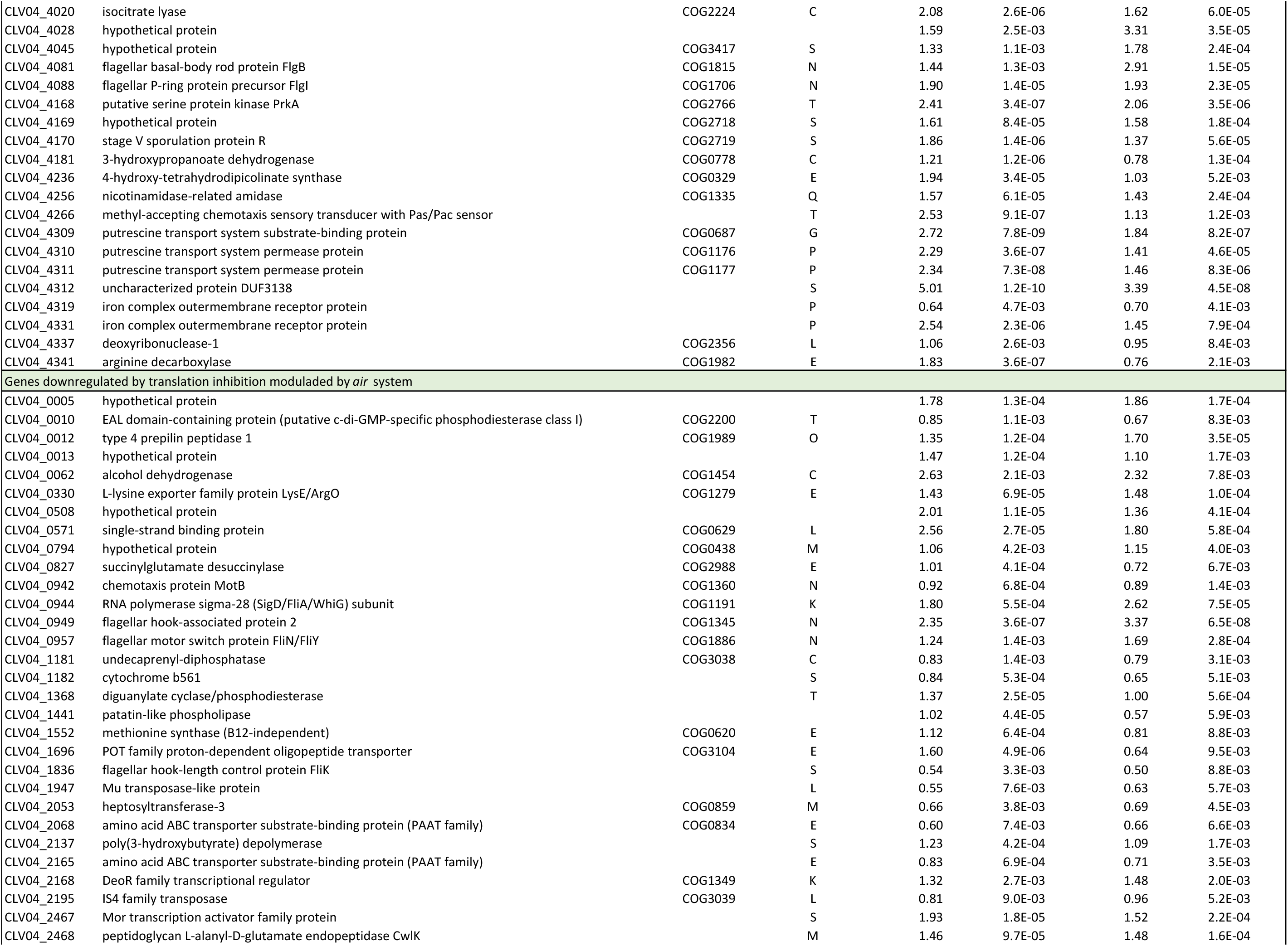
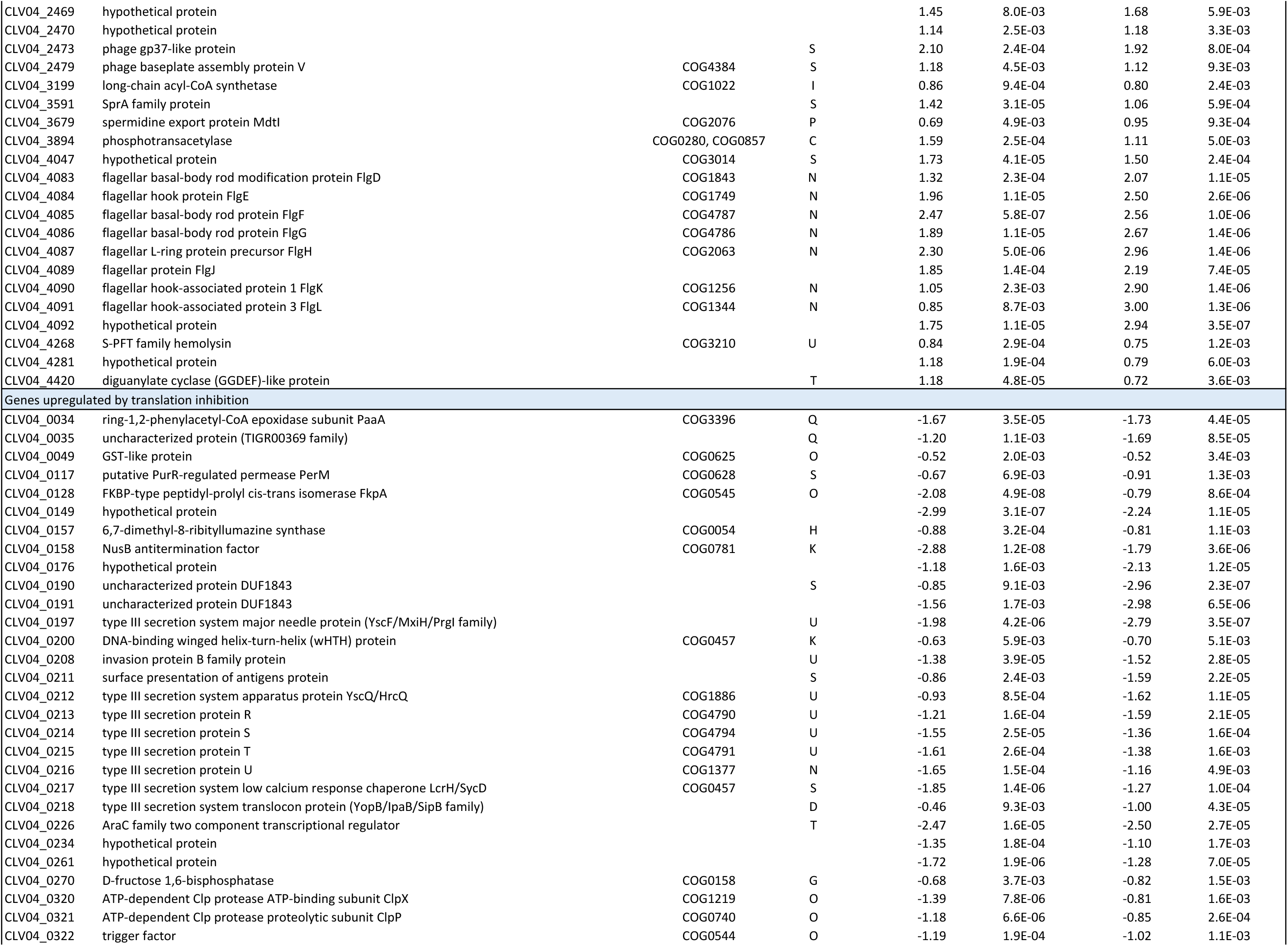
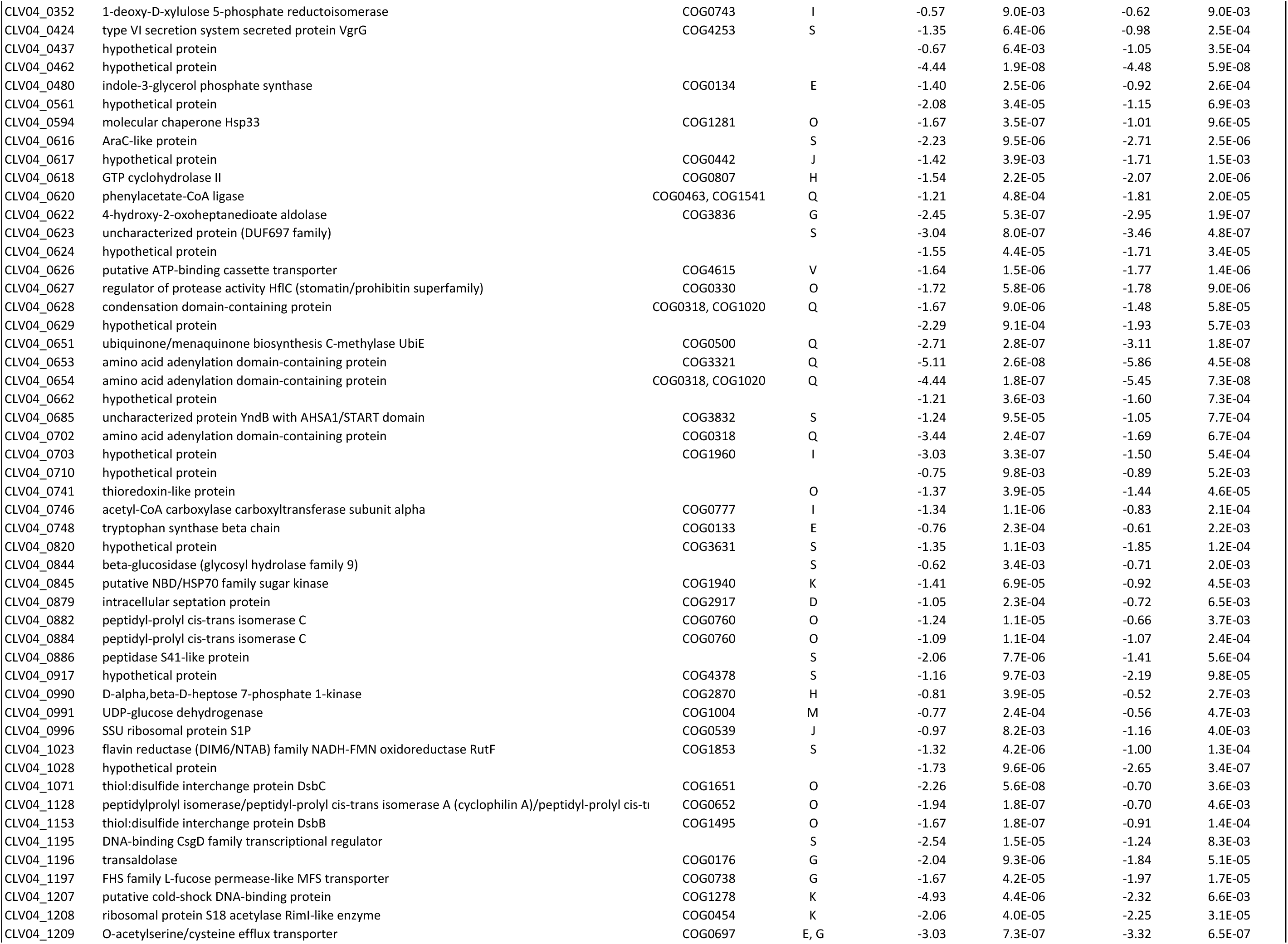
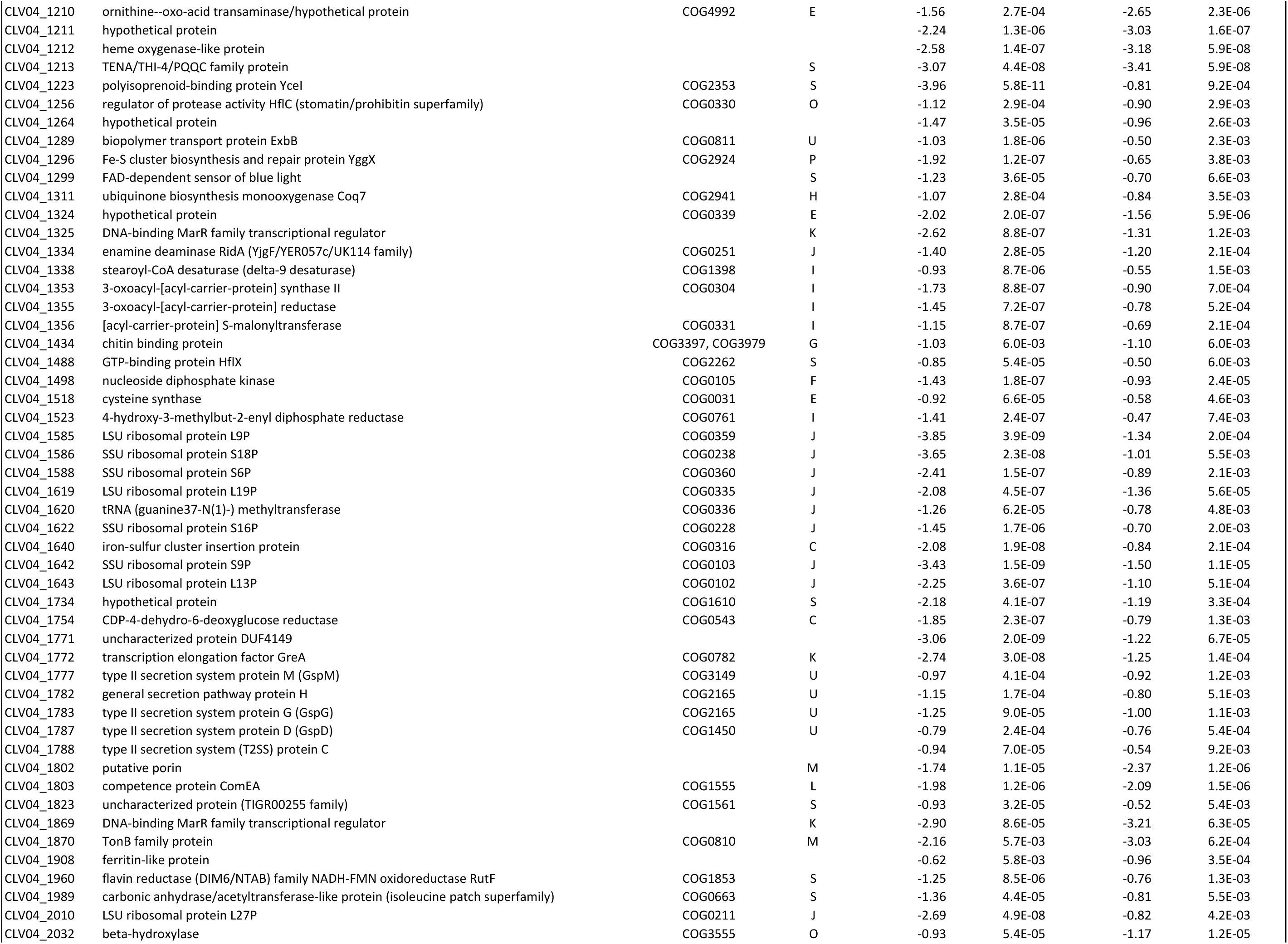
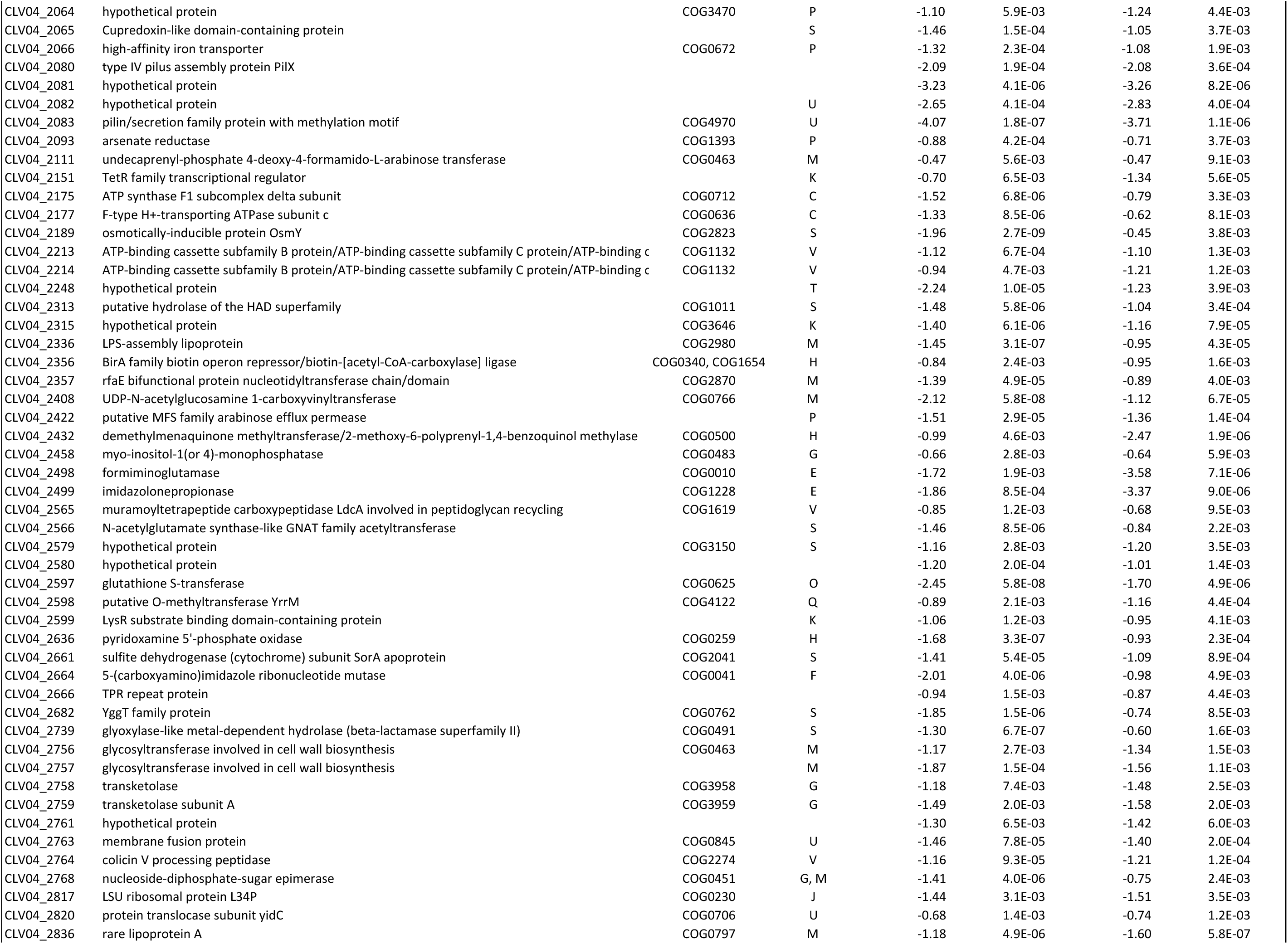
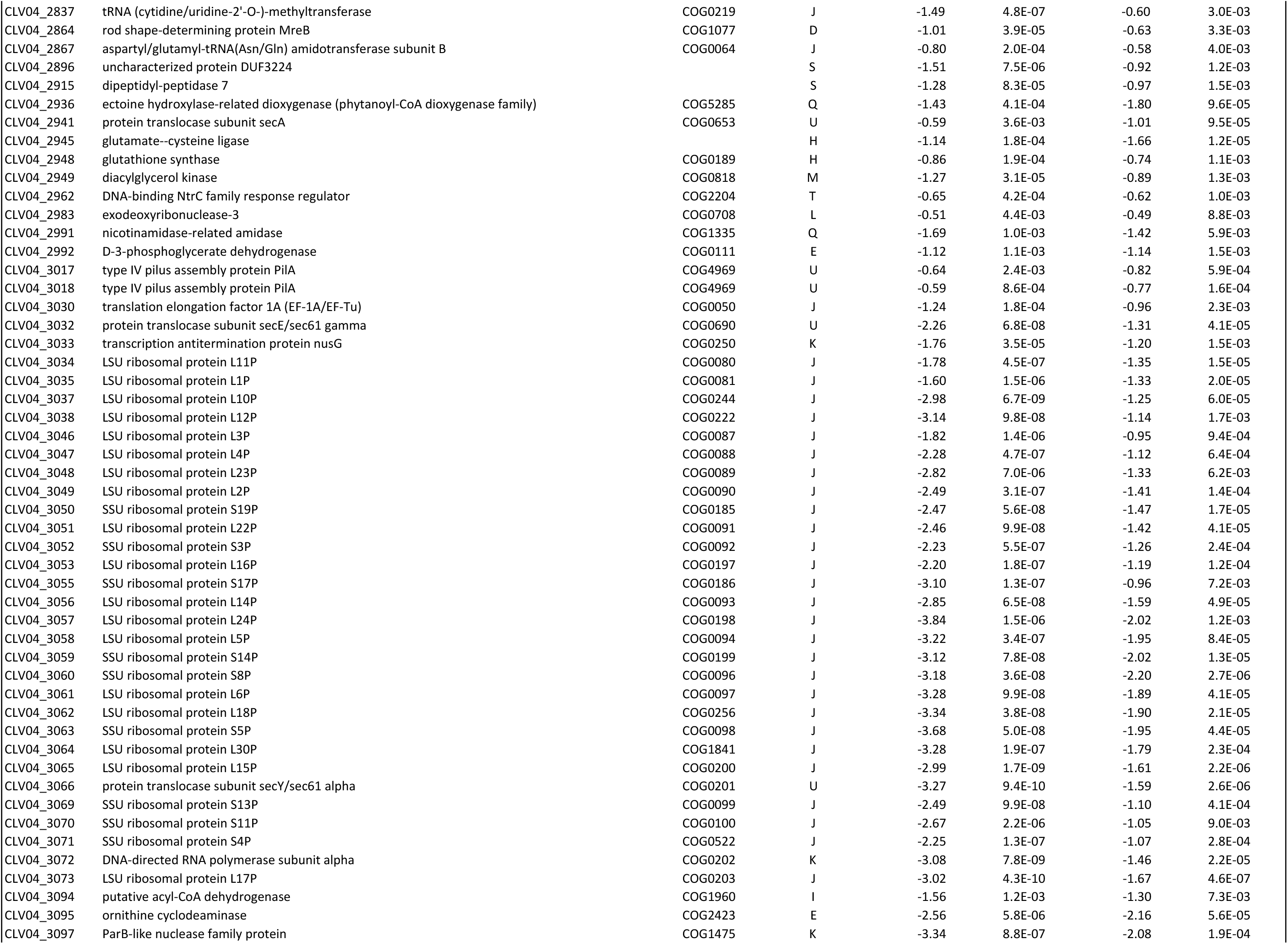
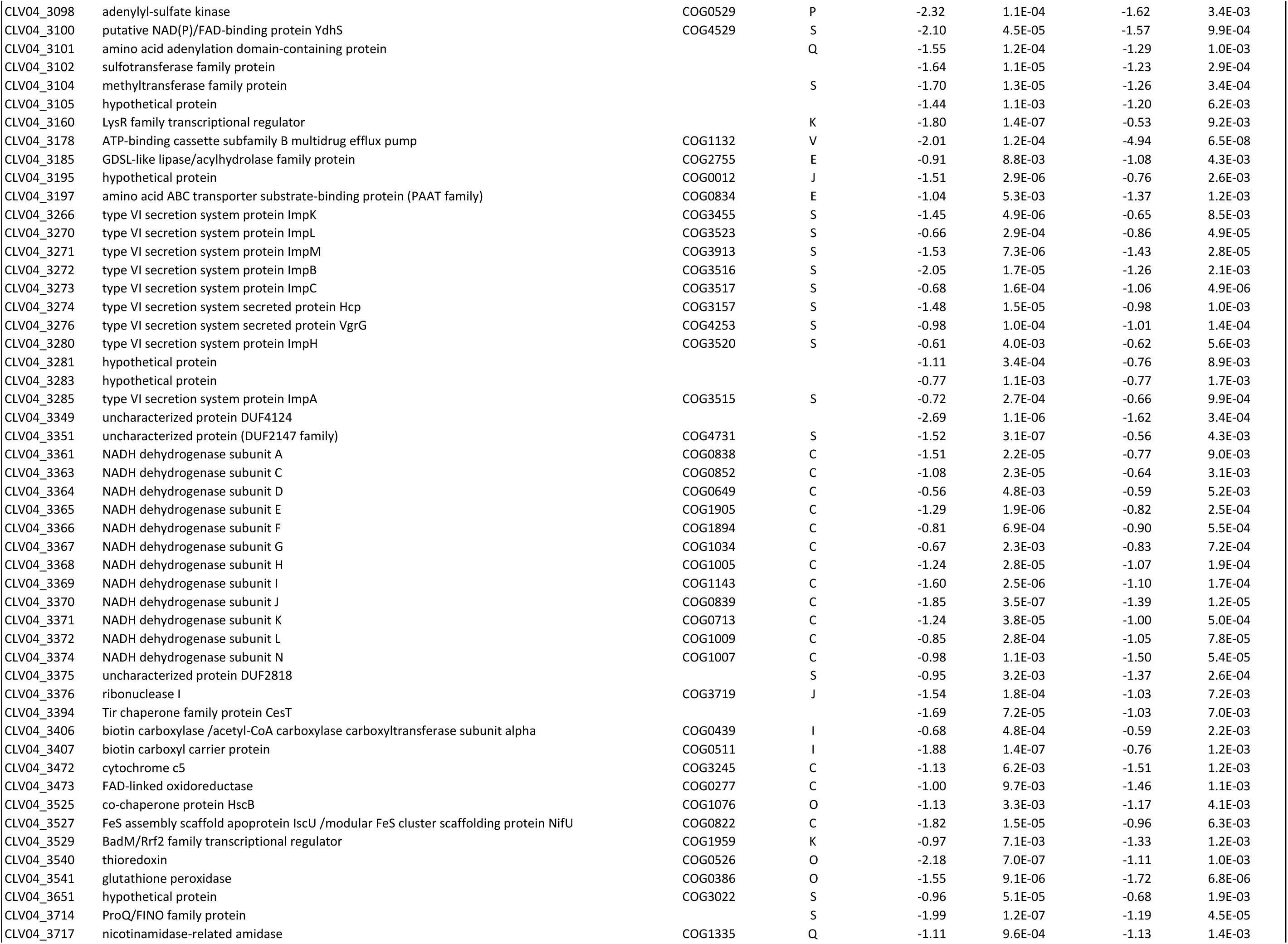
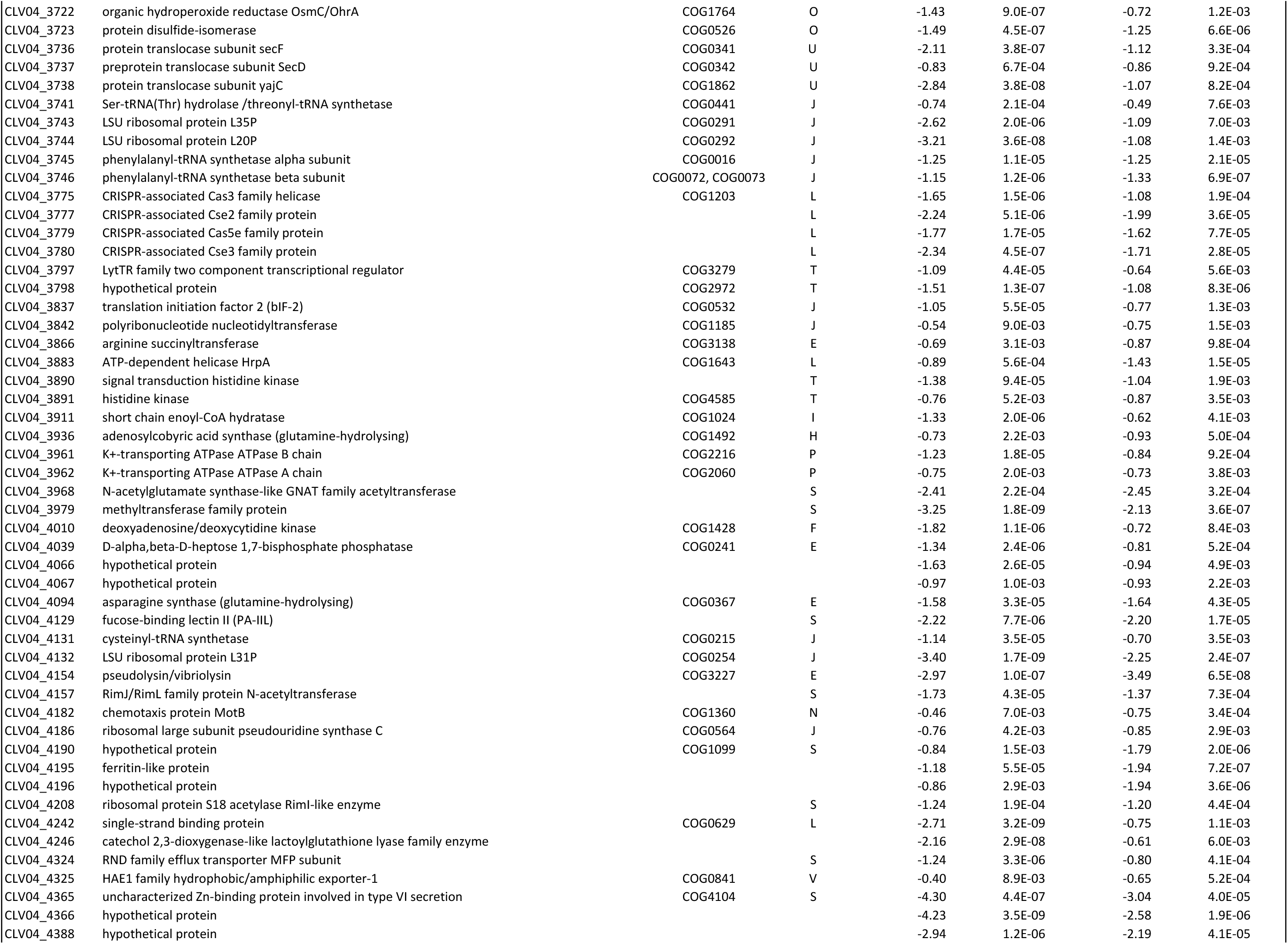
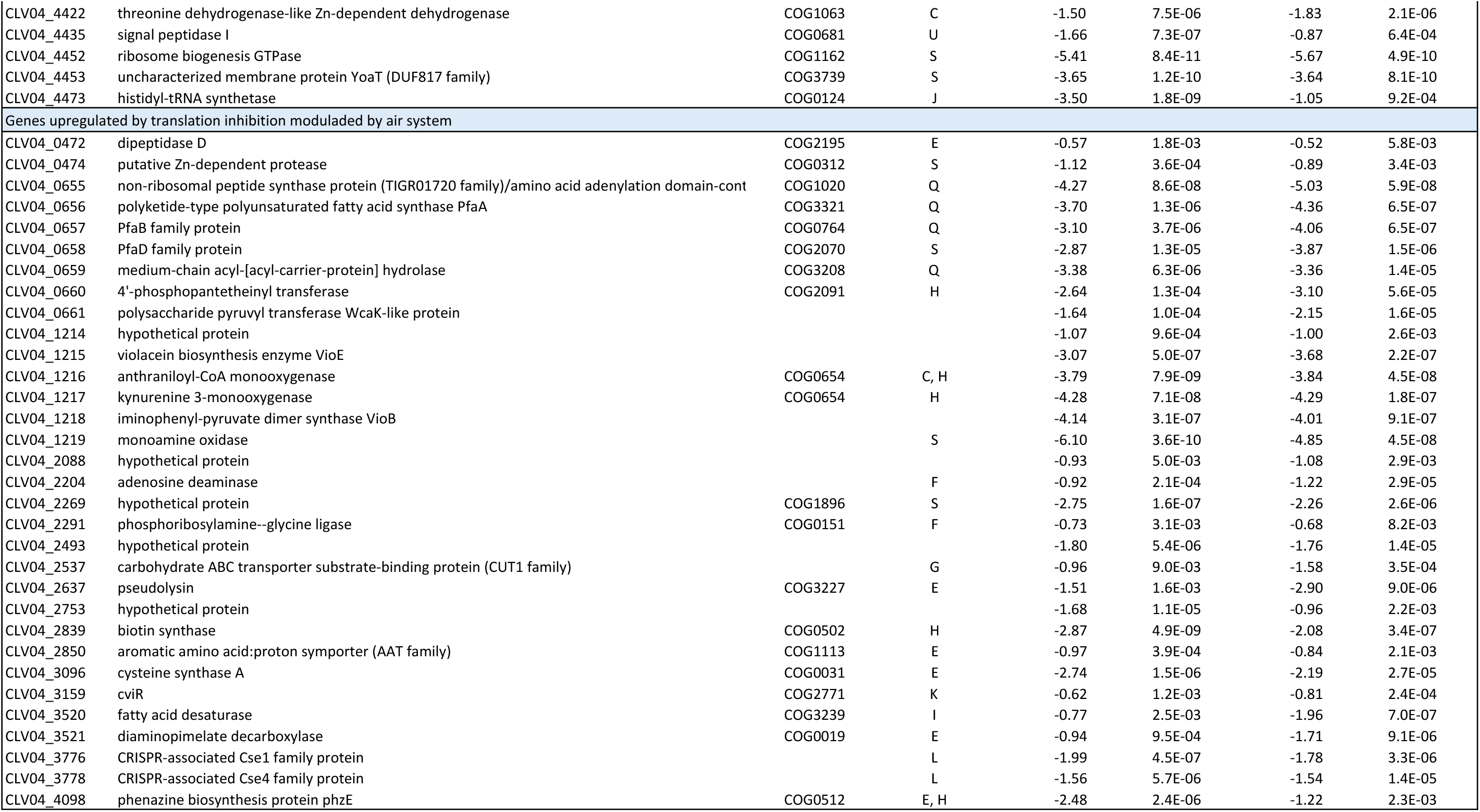
Genes expressed differentially in response to streptomycin and tetracycline.

**Table supplement 2.** Differential gene expression values of the genes identified in the genetic screen.

| Gene ID | Predicted Function | WT |  | WT |  | *airR* |  | *airR* |  |
| --- | --- | --- | --- | --- | --- | --- | --- | --- | --- |
|  |  | No Antibiotic VS Spectinomycin |  | No Antibiotic VS Tetracycline |  | No Antibiotic VS Spectinomycin |  | No Antibiotic VS Tetracycline |  |
|  |  | Log<sub>2</sub>(Fold change) | P Value | Log<sub>2</sub>(Fold change) | P Value | Log<sub>2</sub>(Fold change) | P Value | Log<sub>2</sub>(Fold change) | P Value |
| Conserved response |  |  |  |  |  |  |  |  |  |
| CLV04_2302 | Oxidoreductase molybdopterin binding domain protein | -0.57 | 5.3E-02 | -0.32 | 3.3E-01 | -0.75 | 1.8E-02 | -0.64 | 5.9E-02 |
| CLV04_2301 | Sensor histidine kinase | -0.14 | 5.0E-01 | -0.01 | 9.8E-01 | -0.29 | 1.7E-01 | -0.27 | 2.6E-01 |
| CLV04_3834 | Long chain fatty-acid CoA ligase | 0.02 | 8.9E-01 | **-0.47** | **2.9E-03** | 0.23 | 7.6E-02 | **-0.41** | **7.9E-03** |
| CLV04_1656 | Phosphoenolpyruvate synthase | 0.48 | 2.5E-02 | -0.05 | 8.6E-01 | 0.34 | 1.0E-01 | -0.03 | 9.0E-01 |
| Variable response |  |  |  |  |  |  |  |  |  |
| CLV04_2730 | SAM-dependent methyltransferase | 0.22 | 1.9E-01 | 0.17 | 3.9E-01 | **-0.47** | **9.3E-03** | -0.17 | 3.6E-01 |
| CLV04_2412 | Transcriptional repressor *cdeR* | 0.55 | 1.5E-02 | -0.04 | 8.9E-01 | **-0.56** | **1.1E-02** | 0.25 | 2.9E-01 |
| CLV04_1292 | Phosphoglyceromutase | **0.41** | **8.5E-03** | 0.08 | 6.2E-01 | 0.14 | 3.3E-01 | 0.02 | 8.9E-01 |
| CLV04_2695 | Phosphoglycerate kinase | -0.30 | 2.2E-01 | 0.35 | 1.9E-01 | -0.13 | 6.1E-01 | 0.25 | 3.5E-01 |
| CLV04_1869 | MarR family transcriptional regulator | **-2.90** | **8.6E-05** | **-3.21** | **6.3E-05** | **-2.95** | **6.3E-05** | **-2.51** | **6.2E-04** |
| CLV04_3464 | GntR family transcriptional regulator | 0.47 | 2.71E-02 | 0.44 | 5.4E-02 | -0.19 | 3.2E-01 | 0.07 | 7.6E-01 |
| CLV04_3178 | Multidrug ABC transporter ATP-binding protein | **-2.01** | **1.2E-04** | **-4.94** | **6.5E-08** | **-3.10** | **1.6E-06** | **-4.75** | **1.7E-07** |
| CLV04_1011 | Enoyl-CoA hydratase | **2.93** | **8.6E-08** | **2.53** | **6.5E-07** | **2.86** | **7.1E-08** | **2.21** | **2.3E-06** |

**Table supplement 3.** Genes differentially expressed between *C. violaceum* ATCC31532 WT and *airR* mutant in the absence of antibiotics.

|  |  |  |  | No Antibiotic<br>WT VS *airR* |  |
| --- | --- | --- | --- | --- | --- |
| Gene ID | Predicted Function | COGs ID | COG Category | Log<sub>2</sub>(Fold change) | P Value |
| CLV04_0653 | amino acid adenylation domain-containing protein | COG3321 | Q | 1.84 | 1.1E-02 |
| CLV04_0654 | amino acid adenylation domain-containing protein | COG0318, COG1020 | Q | 2.21 | 1.0E-02 |
| CLV04_0655 | non-ribosomal peptide synthase protein (TIGR01720 family)/amino acid adenylation domain-c | COG1020 | Q | 2.33 | 4.0E-03 |
| CLV04_0656 | polyketide-type polyunsaturated fatty acid synthase PfaA | COG3321 | Q | 2.68 | 3.9E-03 |
| CLV04_0657 | PfaB family protein | COG0764 | Q | 2.88 | 1.8E-03 |
| CLV04_0658 | PfaD family protein | COG2070 | S | 2.83 | 5.1E-03 |
| CLV04_0661 | polysaccharide pyruvyl transferase WcaK-like protein |  |  | 2.08 | 1.8E-03 |
| CLV04_2297 | nicotinamidase-related amidase | COG1335 | Q | -1.38 | 9.8E-03 |
| CLV04_2298 | MarR family transcriptional regulator | COG1846 | K | -1.45 | 1.2E-03 |
| CLV04_2785 | TetR family transcriptional regulator |  | K | 1.23 | 4.3E-03 |
| CLV04_4095 | cytochrome c |  | S | 2.20 | 9.8E-03 |
| CLV04_4096 | p-hydroxybenzoate 3-monooxygenase | COG0654 | Q | 2.79 | 1.2E-03 |
| CLV04_4097 | GMP synthase (glutamine-hydrolysing) | COG0518 | F | 2.36 | 2.2E-03 |
| CLV04_4099 | glutathione S-transferase | COG0625 | O | 1.28 | 4.8E-02 |
| CLV04_4100 | glutamate racemase | COG0796 | M | 1.23 | 2.6E-02 |

**Table supplement 4.**
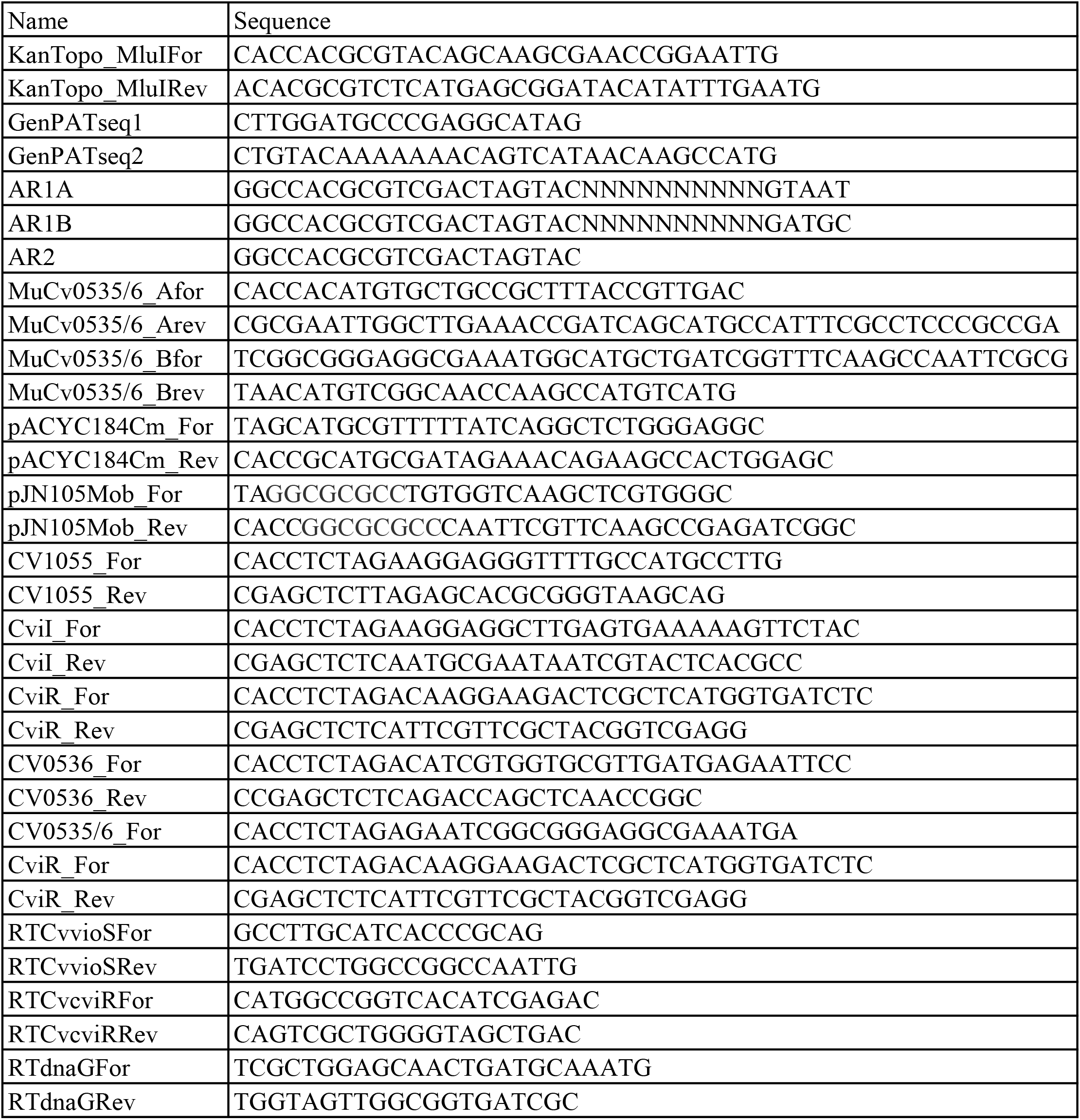
Primers used in this study.

**Table supplement 5.**
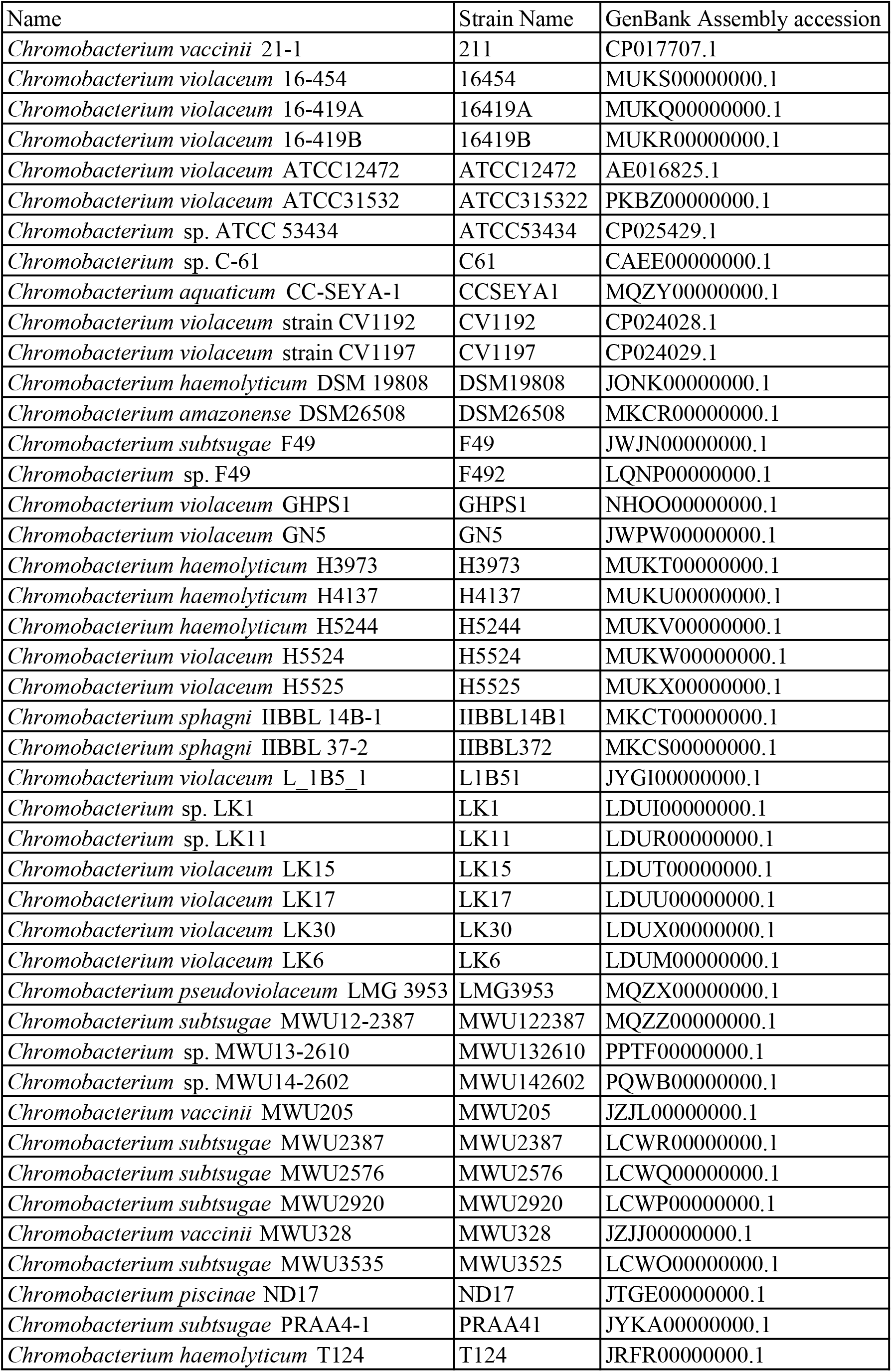
Chromobacterium spp. genomes used for phylogenetic reconstruction.

**Table supplement 6.**
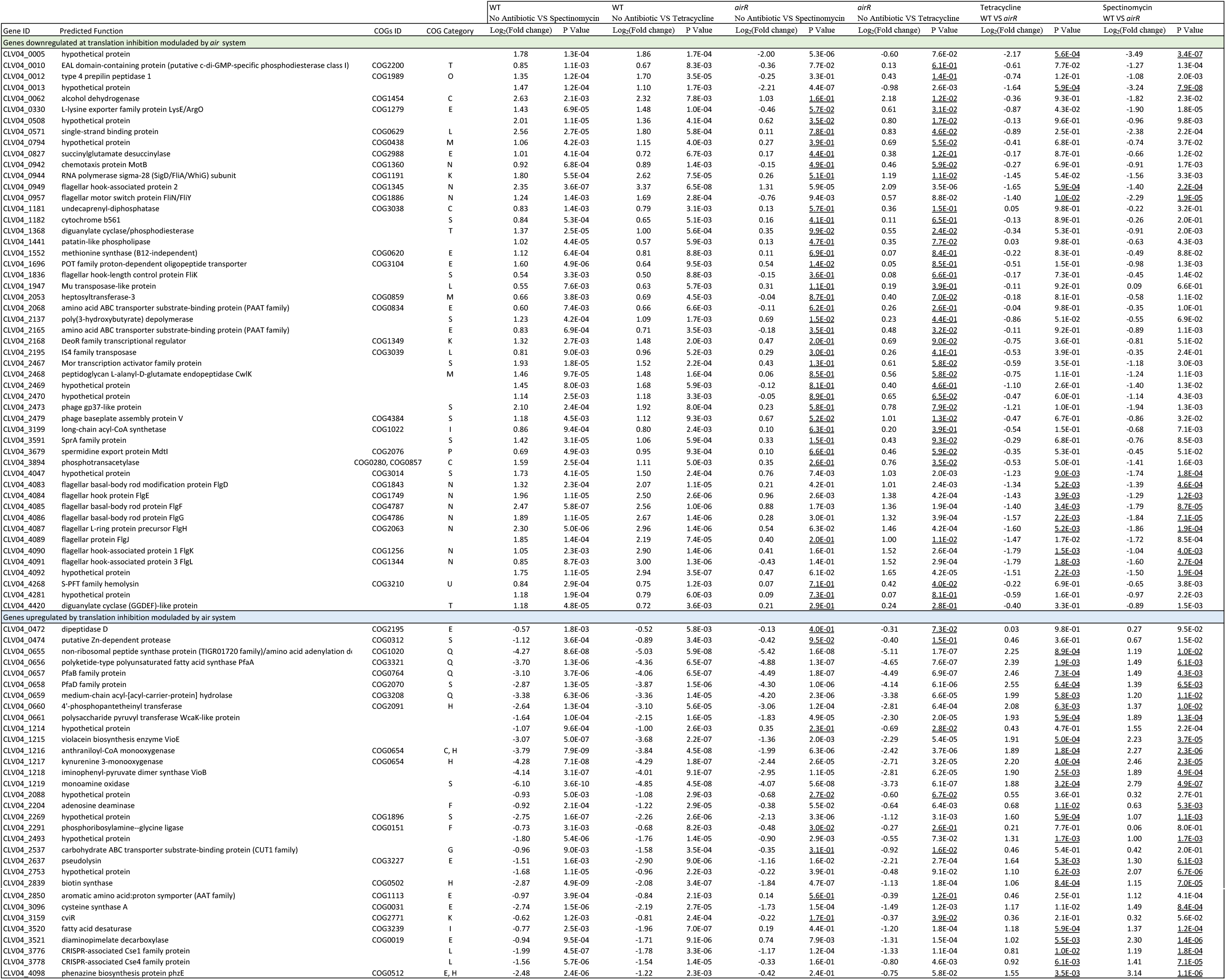
Genes expressed differentially in response to streptomycin and tetracycline that are modulated by *airR*.

